# Trophic flexibility and hydrology structure alpine stream food webs: Implications for a fading cryosphere

**DOI:** 10.1101/2023.02.23.529795

**Authors:** Karen L. Jorgenson, Scott Hotaling, Lusha M. Tronstad, Debra S. Finn, Sarah M. Collins

**Affiliations:** Department of Zoology, University of Wyoming, Laramie, WY, USA; Department of Watershed Sciences, Utah State University, Logan, UT, USA; Wyoming Natural Diversity Database, University of Wyoming, Laramie, WY, USA; Department of Biology, Missouri State University, Springfield, MO, USA

**Keywords:** Alpine stream, food webs, stable isotopes, macroinvertebrates, climate change, *Hydrurus*

## Abstract

Understanding biotic interactions and how they vary across habitats is important for assessing the vulnerability of communities to climate change. Receding glaciers in high mountain areas can lead to the hydrologic homogenization of streams and reduce habitat heterogeneity, which are predicted to drive declines in regional diversity and imperil endemic species. However, little is known about food web structure in alpine stream habitats, particularly among streams fed by different hydrologic sources (e.g., glaciers or snowfields). We used gut content and stable isotope analyses to characterize food web structure of alpine macroinvertebrate communities in streams fed by glaciers, subterranean ice, and seasonal snowpack in the Teton Range, Wyoming, USA. Specifically, we sought to: (1) assess community resource use among streams fed by different hydrologic sources; (2) explore how variability in resource use relates to feeding strategies; and (3) identify which environmental variables influenced resource use within communities. Average taxa diet differed among all hydrologic sources, and food webs in subterranean ice-fed streams were largely supported by the gold alga *Hydrurus*. This finding bolsters a hypothesis that streams fed by subterranean ice may provide key habitat for cold-water species under climate change by maintaining a longer growing season for this high-quality food resource. While a range of environmental variables associated with hydrologic source (e.g., stream temperature) were related to diet composition, hydrologic source categories explained the most variation in diet composition models. Less variable diets within versus among streams suggests high trophic flexibility, which was further supported by high levels of omnivory. This inherent trophic flexibility may bolster alpine stream communities against future changes in resource availability as the mountain cryosphere fades. Ultimately, our results expand understanding of the habitat requirements for imperiled alpine taxa while empowering predictions of their vulnerability under climate change.

## Introduction

Understanding biotic interactions within communities and how they vary across environmental gradients is important for assessing vulnerability to climate change (Blois et al. 2013, HilleRisLambers et al. 2013, Perkins et al. 2010). For example, the trophic level of a species can influence its resilience to flooding or drought, with higher trophic levels having lower stability (Post 2002). Additionally, identifying the basal resources supporting food webs can help identify stoichiometric imbalances that may result from climate change, such as lower growth rate in *Daphnia* fed algae grown under elevated CO_2_ levels (Urabe et al. 2003). Ecosystems above tree line in the alpine (hereafter referred to as alpine ecosystems) are models for studying the effects of climate change due to their rapid warming, habitat loss, low organismal density, and the presence of species living close to their thermal limits (Perkins et al. 2010).

As climate change proceeds, aquatic habitats in the alpine are becoming increasingly homogenized as meltwater sources recede (Hotaling et al. 2017, Birrell et al. 2020). This ongoing habitat homogenization is predicted to decrease regional stream diversity and eliminate range-restricted species (Jacobsen et al. 2012, Giersch et al. 2017). However, recent work suggests that alpine macroinvertebrate diversity may persist longer than previously expected, despite glacier loss (Muhlfeld et al. 2020), perhaps due to cold refugia from subterranean meltwater sources such as rock glaciers (Brighenti et al. 2021, Tronstad et al. 2020). While macroinvertebrate community composition in alpine streams is well-known to vary with hydrologic source (Hieber et al. 2005, Brown et al. 2007, Tronstad et al. 2020), the structure of food webs and resource availability in these ecosystems is less clear (Niedrist & Füreder 2017, Fell et al. 2017), particularly as it relates climate-induced loss of meltwater sources.

Theory suggests that organisms in harsh and unstable environments, such as alpine streams, require trophic flexibility to maintain stability (Saint-Béat et al. 2015, Bartley et al. 2019). Trophic flexibility can increase the resilience of aquatic taxa and communities to environmental perturbation, including rising temperatures, pollution, invasive species, and fire (Lisi et al. 2018, Lewis et al. 2014, Colossi Brustolin et al. 2019, Kortsch et al. 2015). Many aquatic taxa can change their feeding habits in response to seasonal resource pulses, habitat disturbances, or during development (Mihuc 1997, Larson et al. 2018, Lancaster et al. 2005). Flexible feeding strategies may facilitate survival in harsh cold-water environments due to limited and seasonally variable resource availability (Laske et al. 2018, Beaudoin et al. 2001, Zah et al. 2001, Clitherow et al. 2013, Al-Shaer et al. 2015). Macroinvertebrate communities in glaciated catchments can have overlapping trophic niches (Sertić Perić et al. 2021) and can uniformly rely on autochthonous resources despite taxonomic differences (Zah et al. 2001). The inherent dietary flexibility that may be required for life in alpine streams may bolster these communities against climate change and the shifting resources it brings to headwaters.

Dominant hydrologic source is a key driver of habitat makeup and resource availability in high mountain streams (Ren et al. 2019, Slemmons et al. 2013). Thus, a comprehensive understanding of alpine food webs requires insight into how hydrologic sources influence trophic ecology. Important hydrologic sources in alpine streams include glaciers, subterranean ice (e.g., rock glaciers and other “cold rocky landforms”, Brighenti et al. 2021), and snowfields. Glacier-fed streams are phosphorous limited due to high nitrogen and low phosphorous, and have low water temperatures (Slemmons et al. 2013, Ren et al. 2019, Robinson et al. 2002). Additionally, they have low light penetration and high scouring due to large amounts of suspended solids. Rock glacier-fed streams have similar macronutrient and average temperature levels to glacier-fed streams but lower suspended solids (Williams et al. 2007). In contrast, snowmelt streams have high light penetration, more variable temperatures, lower suspended solids, and are more often nitrogen limited (Slemmons et al. 2013, Beck et al. 2021, Warner et al. 2017). These environmental differences can result in snowmelt-fed streams having a larger quantity of autochthonous resources than glacier streams (Uehlinger et al. 2009, Cauvy-Fraunié et al. 2016).

Combining gut content and stable isotope analyses (SIA) is a well-stablished approach for understanding resource use (e.g., Whitledge & Rabeni 1997, Davis et al. 2012). Previous stable isotope investigations of alpine stream food webs have focused on glacier-fed streams (but see Di Cugno & Robinson 2017). Comparing food webs across streams fed by different hydrologic sources provides a larger range of conditions to explore the influence of environmental variables on macroinvertebrate diets and the extent of trophic flexibility, and is therefore necessary to adequately characterize the baseline conditions of alpine stream food webs. It is particularly useful that SIA can distinguish between algal biofilms and the multicellular gold algae *Hydrurus foetidus* (hereafter *Hydrurus*), due to its low carbon isotope signature (Zah et al. 2001, Niedrist and Füreder 2018). *Hydrurus* is widely distributed and abundant during the spring in cold meltwater streams and is likely important for the growth of macroinvertebrates (Niedrist & Füreder 2017, Ward 1994). However, the true scale of this connection, and its geographic scope, is unclear. Additional information from gut content and stable isotope analyses is necessary to assess what resources support the base of alpine food webs and what other trophic connections exist among alpine macroinvertebrate taxa.

Here, we explored how the food web structure of macroinvertebrate communities varied among alpine streams fed by glaciers, subterranean ice, or perennial snowfields. Our primary objectives were: 1) to estimate how the importance of major basal resources (biofilm, plant detritus, and *Hydrurus*) vary with hydrologic sources; (2) to explore how variability in resource use relates to the feeding strategies employed by macroinvertebrates; and (3) to assess relationships between environmental variables (e.g., stream temperature) and resource use within stream communities. We hypothesized that autochthonous resource use would be highest in snowmelt streams, intermediate in subterranean ice streams, and lowest in glacier-fed streams. We also expected that most taxa would display trophic flexibility in response to limited resource availability. Finally, we hypothesized that biofilm importance would be positively correlated with temperature and negatively correlated with total suspended solids, due to their effect on algal growth within biofilms.

## Methods

### Study area

We conducted our study on nine alpine streams in Grand Teton National Park and the adjacent Jedediah Smith Wilderness in Wyoming, USA (Figure 1). The Teton Range is a useful model for our study in several respects. First, glaciers in the Rocky Mountains are melting rapidly (Rice et a. 2018, Edmunds et al. 2012, Hall & Fagre 2003) and glaciers in the Teton Range are already quite small, with the largest—the Teton Glacier—less than 0.22 km^2^ (Edmunds et al. 2012). These glaciers are likely past peak runoff (Chesnokova et al. 2020), which provides insight into the future for other glaciated watersheds. Additionally, the Teton Range hosts glaciers, snowfields, and many rock glaciers (Goff 2019). The range also harbors several species of management concern, including two stoneflies: *Zapada glacier* and *Lednia tetonica* (Hotaling et al. 2019; Green et al. 2022). *Zapada glacier* and a close relative of *L. tetonica* that inhabits similar habitat in Glacier National Park, *Lednia tumana*, were both recently listed as Threatened under the Endangered Species Act (U.S. Fish and Wildlife Service 2019). Finally, the community composition of macroinvertebrates in these streams suggests that subterranean ice-fed streams could act as climate refugia for cold-water taxa (Tronstad et al. 2020).

**Figure 1.**
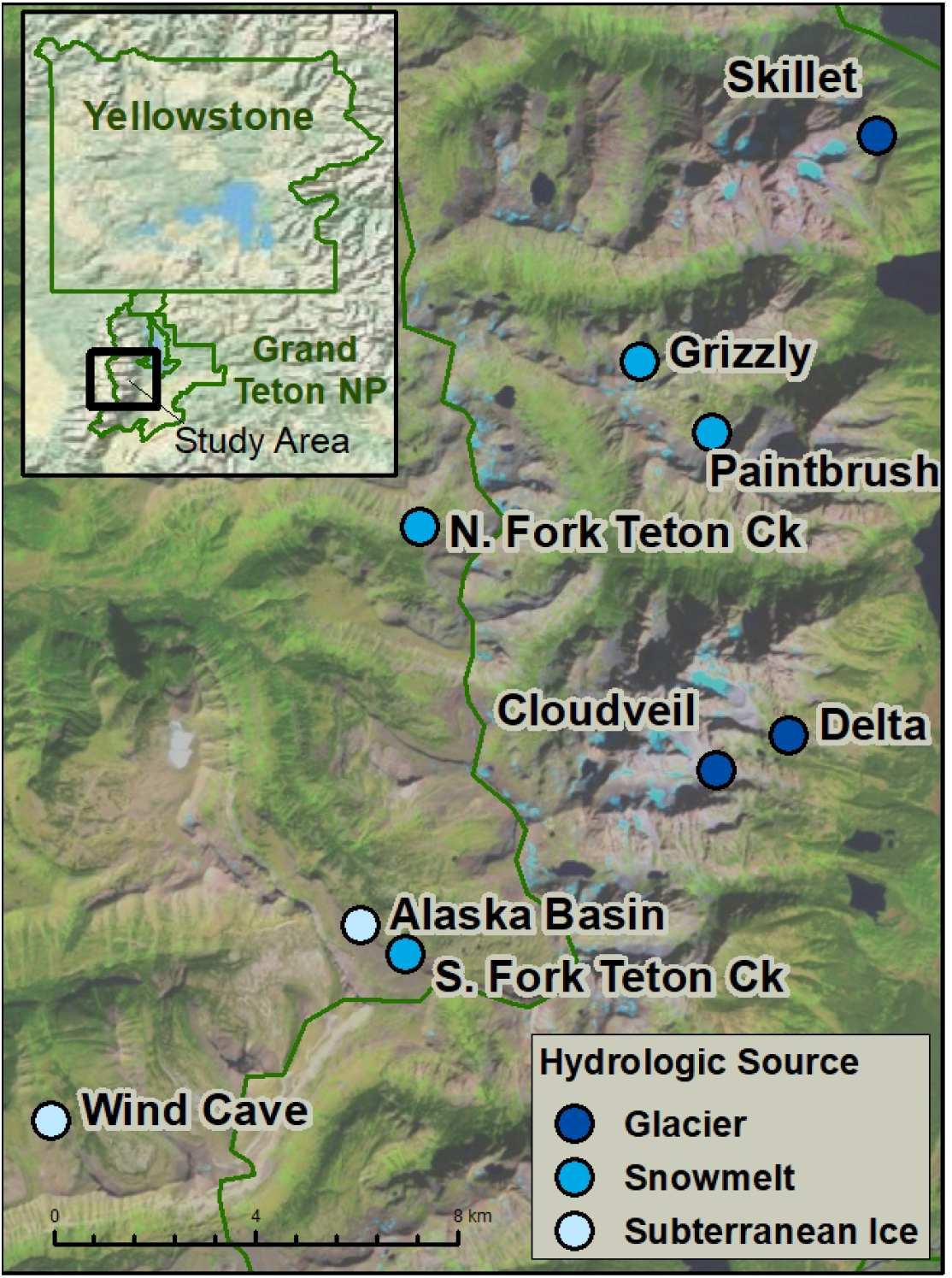
Map of study sites within Grand Teton National Park and the adjacent Jedediah Smith Wilderness in Wyoming, USA.

### Field sampling

Our nine focal streams included two fed by subterranean ice, three fed by surface glaciers, and four fed by perennial snowfields. One of our two subterranean ice-fed streams (Wind Cave) is fed by ice within a cave system and the other (Alaska Basin) is fed by a rock glacier. All sites lacked canopy cover except for one glacier-fed site (Delta), which is at treeline and partially shaded by riparian conifers. All streams were fishless and were sampled near their source and above any lakes to capture as much of the dominant hydrologic source as possible.

From 6–17 August 2020, we collected samples for isotope analysis from each stream. Macroinvertebrates, algae, plant detritus, and any other identifiable food resources were hand-picked from all available stream habitat types (e.g., riffles, pools, etc.). Invertebrates were identified using a hand lens to the lowest taxonomic level (usually genus), which was facilitated by our detailed understanding of what taxa are typically present at these sites (e.g. Tronstad et al. 2020). We degutted invertebrate samples for isotope analysis in the field using forceps and placed the body tissue in snap-cap microcentrifuge tubes by taxa. Invertebrates that were too small to dissect (e.g., midge larvae) were kept in vials for several hours to void their gut contents. At least three replicates of each food item were collected. Macroalgae and stream plant detritus were placed in vials or plastic bags. We also sampled terrestrial plants to ensure the collection of all contributors to stream plant detritus. At least three rocks were scrubbed for biofilm using a plastic brush and stream water. The resulting homogenized slurries were filtered onto pre-weighed Whatman GF/F glass-fiber filters (0.45-μm pore size) using a 60-ml syringe before being stored in tinfoil. We kept isotope samples cool on overnight backpacking trips by burying them in snow or submerging them within a stream., and they were frozen within 36 hours. Additional samples were collected during 20-28 July 2021 to supplement samples of *Hydrurus* or biofilm.

We also collected macroinvertebrate samples for visual gut content analysis in August 2020 and July 2021. Individuals of abundant taxa were preserved in 95% ethanol (2020) or Kahle’s solution (2021) at the field site. We switched to Kahle’s solution (Rosi-Marshall et al. 2016) for easier dissection and observed that both methods seemed to preserve the gut contents well. Samples of each food resource were also preserved for reference materials.

We collected quantitative benthic invertebrate samples and measured a suite of environmental variables at each site in August 2020. Invertebrate abundance and diversity were calculated from three Surber samples taken along the length of the reach. Dissolved oxygen (DO), specific conductivity (SPC), and pH were measured using a Yellow Springs Instrument (YSI) Professional Plus multiparameter sonde that was calibrated at the trailhead (SPC and pH) or at the site (DO). Recorded volumes of water were filtered onto ashed and pre-weighed 25 mm glass-fiber filters to measure total suspended solids (TSS). We collected filtered water samples to be analyzed for anions (nitrate, sulfate, chloride, and fluoride). Temperature loggers (HOBO Pro v2, Onset Computer Corporation) were secured to rocks on the streambed to record stream temperature every hour year-round. Elevation, slope, and aspect were compiled from measurements from previous sampling years.

### Laboratory analyses

We analyzed macroinvertebrates and food resources for carbon (C) and nitrogen (N) isotopes. We dried isotope samples at 60 °C for at least 72 hours. Invertebrate samples were homogenized in microcentrifuge tubes using pestles and were tinned whole or homogenized individually or in groups to achieve adequate sample mass. We weighed biofilm samples on filters, calculated biofilm mass by subtracting the initial filter weight, and then tinned the filters. Plant samples and detritus were homogenized with a mortar and pestle, and *Hydrurus* samples were homogenized with a pestle in a microcentrifuge tube before tinning. Samples were analyzed at the University of Wyoming Stable Isotope Facility for *δ*^13^C and *δ*^15^N using a Carlo Erba 1110 Elemental Analyzer coupled to a Thermo Delta V isotope-ratio mass spectrometer (IRMS).

Gut contents were visually quantified following Rosi-Marshall et al. (2016) with the following modifications. The foregut or front third of the invertebrate guts were mounted onto slides. We photographed two random transects across each slide at 200x magnification using a compound microscope. We assigned particles to the following categories: diatoms, *Hydrurus*, plant detritus, animal material, or “other.” Particles assigned to “other” made very small contributions to the total area measured and included other filamentous algae, fungal hyphae, and moss rhizoids. The area of each category was measured across multiple photos and then divided by the total area of those photos to calculate proportional cover. For abundant particles, 3-5 photos were used, while rare particles were measured across both transects (∼50 photos). We calculated percent composition of each diet item by dividing proportional cover by the sum of proportional covers for all categories and multiplying by 100. The percent assimilated from each diet item was calculated from the percent composition with assimilation factors of 0.3 for diatoms, 0.1 for plant detritus, 0.3 for filamentous algae (which we used for *Hydrurus*), and 0.7 for animal material (Benke & Wallace 1980, Cross et al. 2013).

We analyzed environmental variables to assess differences in the physical and chemical habitat among streams. We dried TSS samples for at least 72 hours at 60 °C and calculated TSS as the difference between the sample and filter dry weight and the initial filter weight. Filtered water samples were measured for anion concentrations using an Ion Chromatograph (Thermo Scientific Dionex Dual Integrion RFIC). For a few data points that were below the detection limit, we used values halfway between zero and the detection limit (12.27 μg/L for nitrate and 21.48 μg/L for fluoride). We calculated the mean (T_mean_) and max (T_max_) water temperatures for the two-week period prior to our 2020 sampling. For Wind Cave, temperature data from 2019 were used due to a lost temperature logger in 2020, which was likely similar to 2020 because the stream temperature at this site is very stable across years (Tronstad et al. 2020, Hotaling et al. 2019). The aspect data were transformed using the formula A’ = cos(45 – A) + 1, where A is the aspect in degrees (Beers et al. 1966).

Macroinvertebrates from the Surber samples were used to calculate diversity and biomass. We measured the lengths of up to 40 individuals of each taxon from each sample. We calculated the biomass of each taxon using taxon-specific length-mass regressions from Benke et al. (1999). Invertebrate diversity was estimated for Shannon and Simpson diversity indices (Shannon 1948, Simpson 1949).

### Statistical analyses

All statistical analyses and generation of figures were conducted in R (v4.0.2, R Core Team 2020). We used the Bayesian mixing-model package ‘MixSIAR’ (Stock et al. 2018) to estimate the proportional contribution of resources to invertebrate tissues and establish food web linkages. Separate models were run for each site. Taxon was included as a fixed effect in all models and the default generalist priors were used. We included both process and residual error in the models to incorporate variation in taxa consuming material from different locations on the source distributions and consumer variation not explained by source isotopic value (e.g., individual assimilation, metabolism), respectively (Stock & Semmens 2016). We used Trophic Enrichment Factors (TEFs) of 0.4 ± 1.4 ‰ for *δ*^13^C (Post 2002) and 1.4 ± 1.4 ‰ for δ^15^N (Bunn et al. 2013). TEF selection is discussed further in Appendix 1. Model convergence was assessed by the Gelman diagnostic, the Geweke test, and visual assessment of trace plots.

The food resources included in our model were biofilm, coarse particulate organic matter (CPOM), and *Hydrurus*. Plant and within-stream plant detritus samples were combined into CPOM due to overlapping isotope signatures. Items that were collected as potential food resources but were not observed in the guts of any taxa were excluded from the model (e.g., moss). We included an additional resource for two sites. First, we included a filamentous alga collected at South Fork Teton Creek because an end member representing algae was missing for this site, and a few strands of this alga were observed in some invertebrate guts at this site. Second, we included small mammal fecal pellets collected at Grizzly that provided a missing end member and had caddisfly larvae congregated around them at the site. Sample sizes for different resources was supplemented by collections in July 2021 for *Hydrurus* at Alaska Basin, Skillet and Delta, and biofilm at Skillet. Comparison of samples from different sampling events did not show clear seasonal or annual differences. Source sample sizes ranged from 2 to 14 samples.

We decided which taxa and individuals to include in our primary consumer models using their trophic position (TP) and gut content analysis data. We calculated the trophic position of each individual from the δ^15^N values (δ^15^N_consumer_) using a one-baseline model: TP = 2 + (δ^15^N_consumer_ – δ^15^N_base_)/Δδ15N (Post 2002). We used the TEF (Δ δ^15^N) of 3.4 ‰ from Post (2002) to account for predators. Midges (which did not include any predatory Tanypodinae), or *Allomyia* caddisflies if midges were insufficiently abundant, were selected as the baseline taxon (δ^15^N_base_). Taxa were categorized as predators if animal material was observed in their guts at any site, and individuals of predatory taxa were excluded from the mixing models if their trophic position was calculated to be greater than 2.5.

We used the diet estimates from our isotope mixing models in additional analyses aimed to identify cross-site patterns. Prior to these, we combined the model estimates for filamentous algae and biofilm at South Fork Teton Creek, and CPOM and small mammal feces at Grizzly. We then calculated the average diet composition for each site to avoid pseudoreplication in our analyses. We also used biomass estimates and modeled diet proportions to calculate the invertebrate biomass supported by each food resource in each stream as an alternative way to assess community resource use.

We performed principle component analysis (PCA) to visualize diet variation among sites and environmental variation (e.g., source) using the ‘vegan’ package (Oksanen et al. 2022). We calculated multivariate metric variances (Pawlowsky-Glahn & Egozcue 2001) using the package ‘compositions’ (Van den Boogaart & Tolosana-Delgado 2013) to estimate within-site and among-site variation in the diet compositions. We ran an ANOVA to test whether the within-site variance in diet compositions of taxa significantly differed among hydrologic sources. To evaluate whether trophic diversity related to community diversity, we ran linear models using the lm function to test if the within-site variation in diet compositions increased with increasing Shannon or Simpson diversity.

We used the package ‘DirichletReg’ (Maier 2013) to preform multivariate regression of relationships between diet composition and environmental variables. This package uses a Dirichlet distribution to model compositional data and we used the common parameterization. We modeled the diet composition with the hydrologic source, the first and second principle components of the environmental PCA, and each separate environmental variable. We also ran models with multiple environmental variables as explanatory variables and with environmental variables combined with hydrologic source. However, with our sample size we were not able to combine all of the environmental variables in a single model. We calculated corrected Akaike information criterion (AICc) to determine which variables best modeled our observed diet proportions. Using the same methods, we also modeled the percent biomass at each site that was supported by each resource.

## Results

### Food web structure

Gut content analysis revealed high omnivory in many taxa. While functional feeding group assignments rely on the idea that each taxa has a consistent food source in all environments, our gut content and stable isotope analyses suggest that the alpine macroinvertebrate diets in our study vary across sites. We found omnivory that deviated from the functional feeding groups described by Merrit et al. (2008) for these taxa (Appendix 2, Table S1). Predatory feeding was common, with over half of all taxa (*Megarcys, Rhyacophila, Sweltsa, Prosimuliium*, Ameletidae, *Drunella, Homophylax, Epeorus*, and *Cinygmula*) had midge larva in their guts (Figure 2). Two samples of *L. tetonica* at one site (Delta) had especially high trophic positions (Appendix 2, Figure S1), although we did not see identifiable animal material in their guts. The only observations of wood—cross sections of tracheids—in any digestive tract were from *L. tetonica* and *Zapada* sp. at one site (Delta).

**Figure 2.**
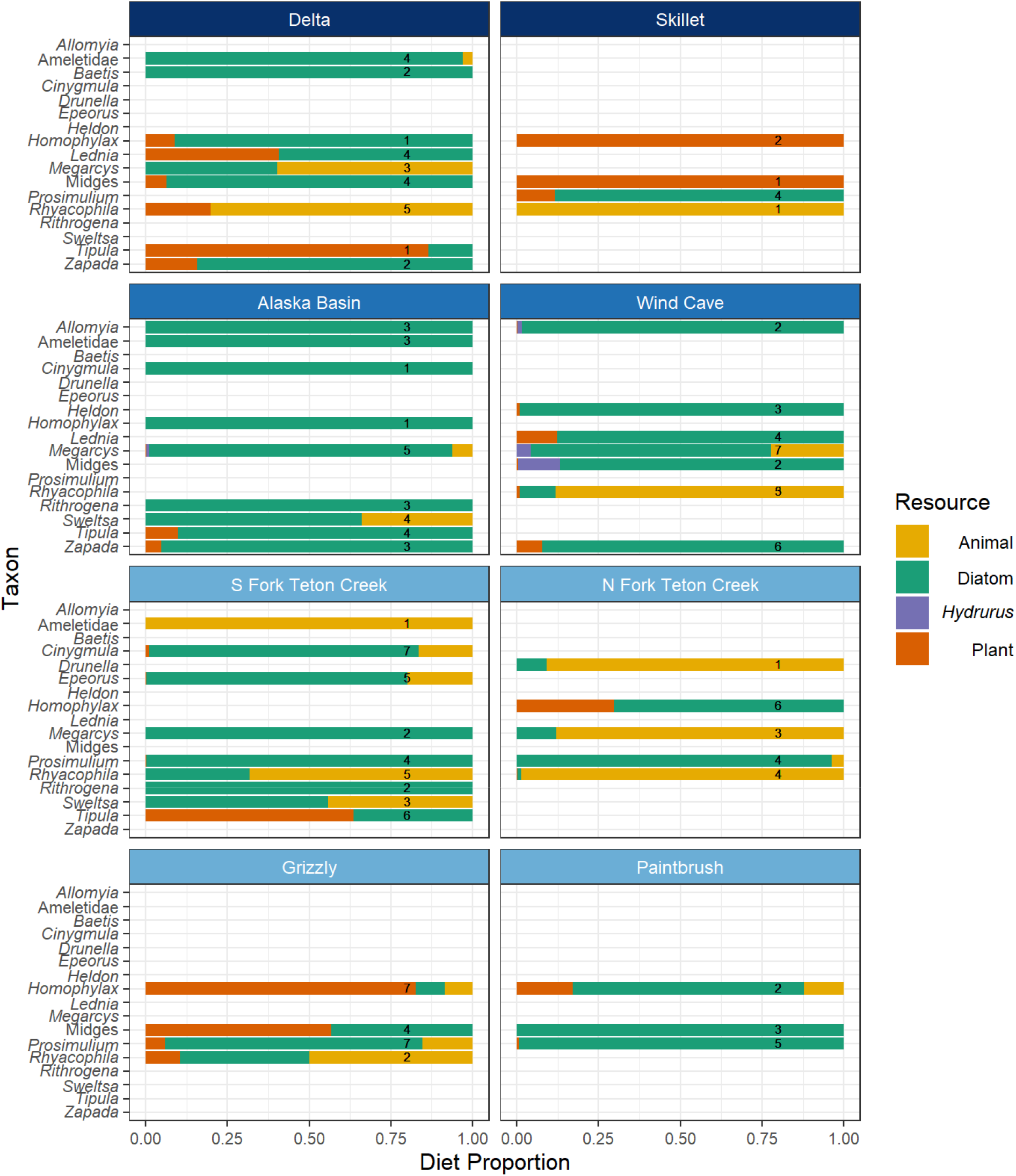
Diet composition for each taxon estimated using gut content analysis. The numbers on each bar represent samples size (multiple individuals were pooled for most samples). Site names are colored by hydrologic source (dark blue = subterranean ice, medium blue = glacier, light blue = snowmelt). Cloudveil is not shown because no identifiable particles were recovered from specimen digestive tracts. This is likely due to low diatom abundance at this site. Very small proportions of a filamentous alga were observed in the guts of *Tipula* and *Prosimulium* at South Fork Teton Creek.

Our results from gut content analysis informed our decisions about which taxa or individual samples to include in the isotope mixing models as primary consumers. It is possible that the fragments of exoskeleton found in the guts of *Homophylax, Epeorus*, and *Cinygmula* were devoid of digestible animal material before they were consumed, and we therefore included all individuals of these taxa in our models. All other taxa observed to contain animal material had some samples above and below the trophic cutoff (Appendix 2, Figure S1). In these cases, only the samples below the cutoff were included. *L. tetonica* over the trophic cutoff were also excluded at Delta. No Turbellaria were included in our model as they were clearly secondary predators in these streams. The food resources had varying isotopic signatures among sites which necessitated the use of site-specific source isotope values in the mixing models. The site δ^13^C mean values of biofilm (−28.9 to −11.6‰) and *Hydrurus* (−34.6 to −22.0‰) were variable across sites, while CPOM was more consistent (−29.3 to −26.3‰). Plots of isotope values at each site are included in the Supplementary Materials (Appendix 2, Figures S2-S10).

Diet proportions for individual taxa varied widely among sites (Figure 3). PCA showed that our sites grouped into glacier, snowmelt, and subterranean ice streams by diet compositions (Figure 4a). Site mean diet compositions were significantly different among streams fed by different hydrologic sources (Figure 5). The diet proportions of biofilm, CPOM and *Hydrurus* were significantly (α = 0.05) different between subterranean ice and both glacier (biofilm: coefficient = −2.0, p-value = 0.030; CPOM: -coefficient = 4.9, p-value < 0.001; *Hydrurus:* coefficient = −6.3, p-value < 0.001) and snowmelt streams (biofilm: coefficient: *-2*.*4, p-value = 0*.*007*; CPOM: coefficient = −4.1852, p-value < 0.001; *Hydrurus:* coefficient = −5.1, p-value < 0.001), but glacier and snowmelt streams did not differ significantly in diet proportion*s* (model results in Appendix 2, Table S2). We observed some differences in the diet compositions of individual taxa between hydrologic sources. For *Zapada*, the proportions of all resources were significantly different between glacier and subterranean ice-fed streams (biofilm: coefficient: *-2*.*6, p-value = 0*.*009*, CPOM: coefficient = 2.2, p-value = 0.032, *Hydrurus:* coefficient = 2.8, p-value = 0.006). The macroinvertebrate biomass supported by each resource followed similar trends due to the low diet variability between taxa within each site (Figure 5b). Glacier and snowmelt-fed streams did not have significantly different biomass contributions, but they both differed from subterranean ice-fed streams in the proportions of biomass supported by CPOM (glacier: coefficient = −2.9, p-value = 0.002; snowmelt: coefficient = −1.9, p-value = 0.030) and *Hydrurus (*glacier: coefficient = −2.8, p-value = 0.003; snowmelt: coefficient = −4.0, p-value < 0.001).

**Figure 3.**
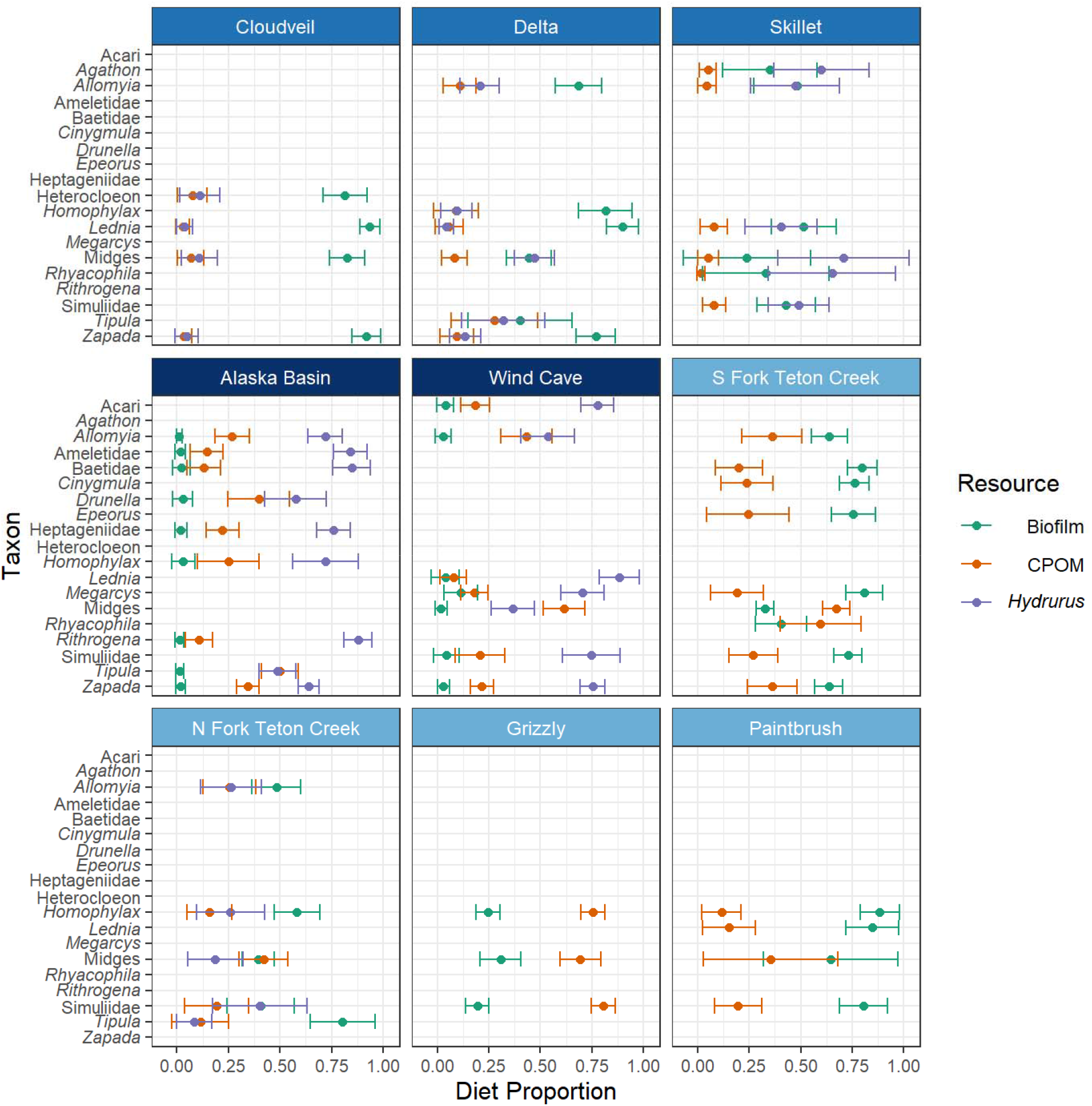
Diet proportions for taxa across sites estimated using isotope mixing models. Error bars represent standard deviation, and the site names are colored by hydrologic source (dark blue = subterranean ice, medium blue = glacier, light blue = snowmelt). Model estimates were combined for the filamentous alga and biofilm at South Fork Teton Creek, and small mammal feces and CPOM at Grizzly.

**Figure 4.**
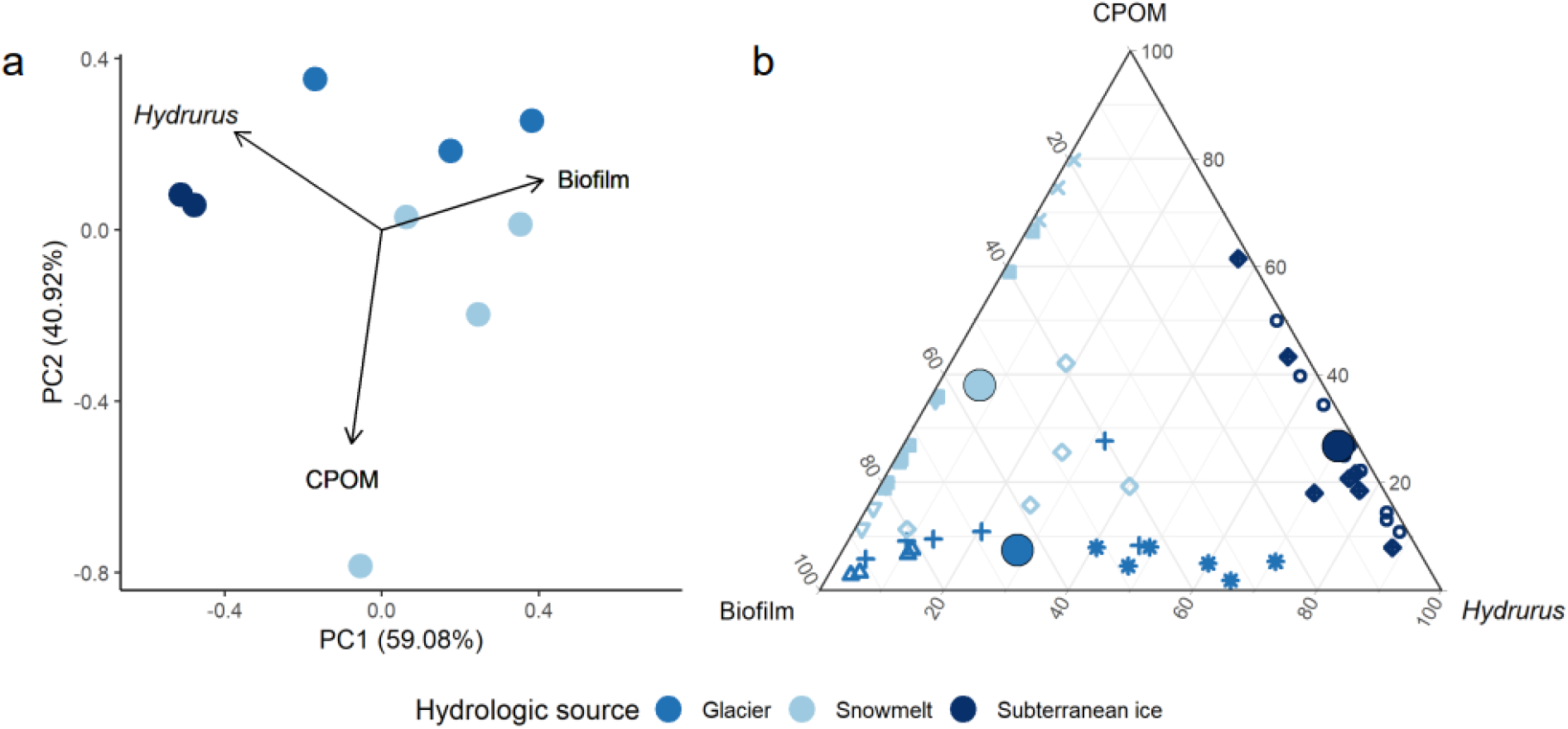
Sites have low within-site variation and are clustered by hydrologic source. (a) Principle component analyses showing diet variation among sites with different hydrologic sources. (b) Ternary plot of taxa diet compositions. Shapes represent different stream sites and large circles depict the mean of each hydrologic source.

**Figure 5.**
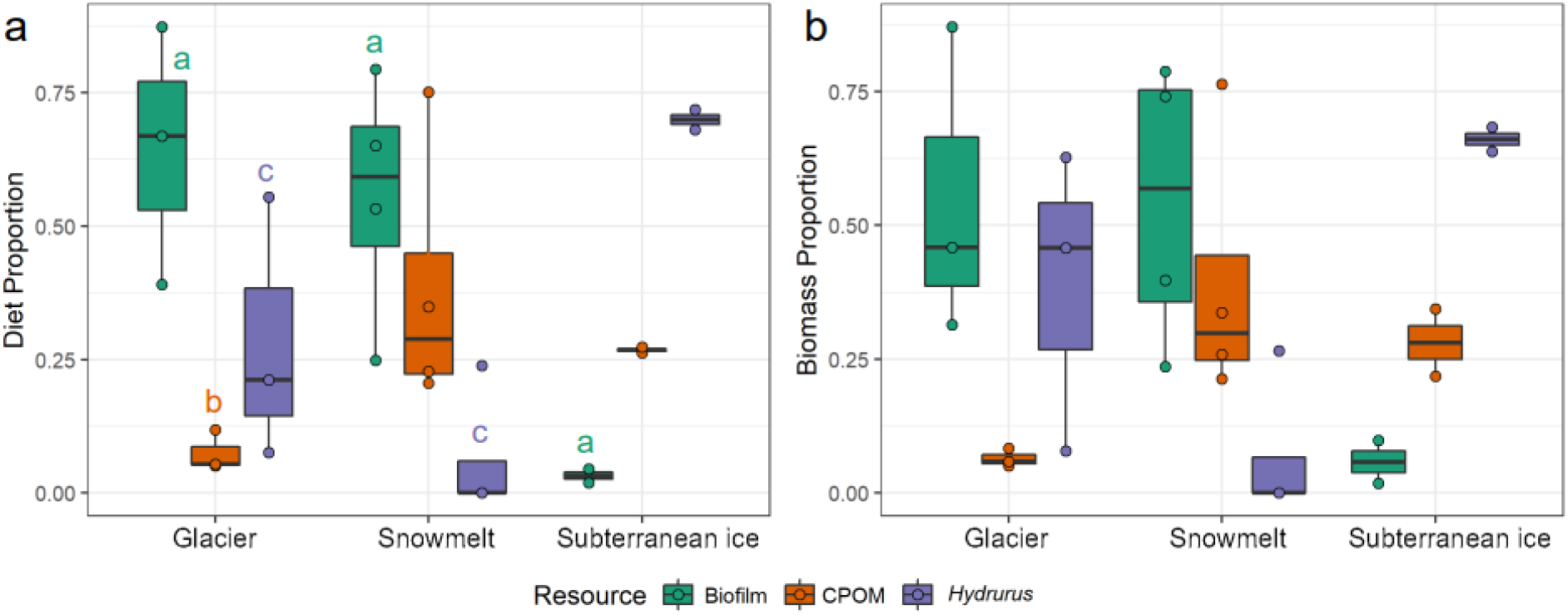
(a) Boxplot of mean site diet proportions of each food resource grouped by hydrologic source. Points represent site means and letters indicate resource proportions that did not differ significantly. Diet compositions were significantly different among all hydrologic sources. (b) Boxplot of the proportion of macroinvertebrate biomass supported by each resource within different hydrologic source categories. Biomass composition was significantly different between subterranean ice and both snowmelt and glacier-fed streams. Points represent unique sites.

The among-site multivariate metric variation was 0.23 and the mean within-site variation was 0.04. Within-site variation did not differ significantly between hydrologic sources, and among-site variation increased from subterranean ice (0.001), to glacier (0.121), to snowmelt (0.132) fed streams. Resource use by *L. tetonica* and *Zapada* ranged from 88% *Hydrurus* and 4% biofilm to 4% *Hydrurus* and 94% biofilm, and 75% *Hydrurus* and 3% biofilm to 5% *Hydrurus* and 92% biofilm, respectively. The mean diet proportions of CPOM across all sites by *L. tetonica* and *Zapada* were 0.08 ± 0.05 and 0.21 ± 0.15. The within-site variation did not have a significant linear relationship with the Shannon or Simpson diversity indices.

### Influence of environmental variables

Site characteristics varied among streams (Table 1). Principle component analysis (PCA) showed that our sites grouped into glacier, snowmelt, and subterranean ice streams by environmental variables (Figure 6). The first principle component (PC1) was primarily influenced by mean temperature (T_mean_), maximum temperature (T_max_), DO, chloride and nitrate. The second principle component (PC2) was primarily influenced by SPC, pH, fluoride, and sulfate.

**Table 1.**
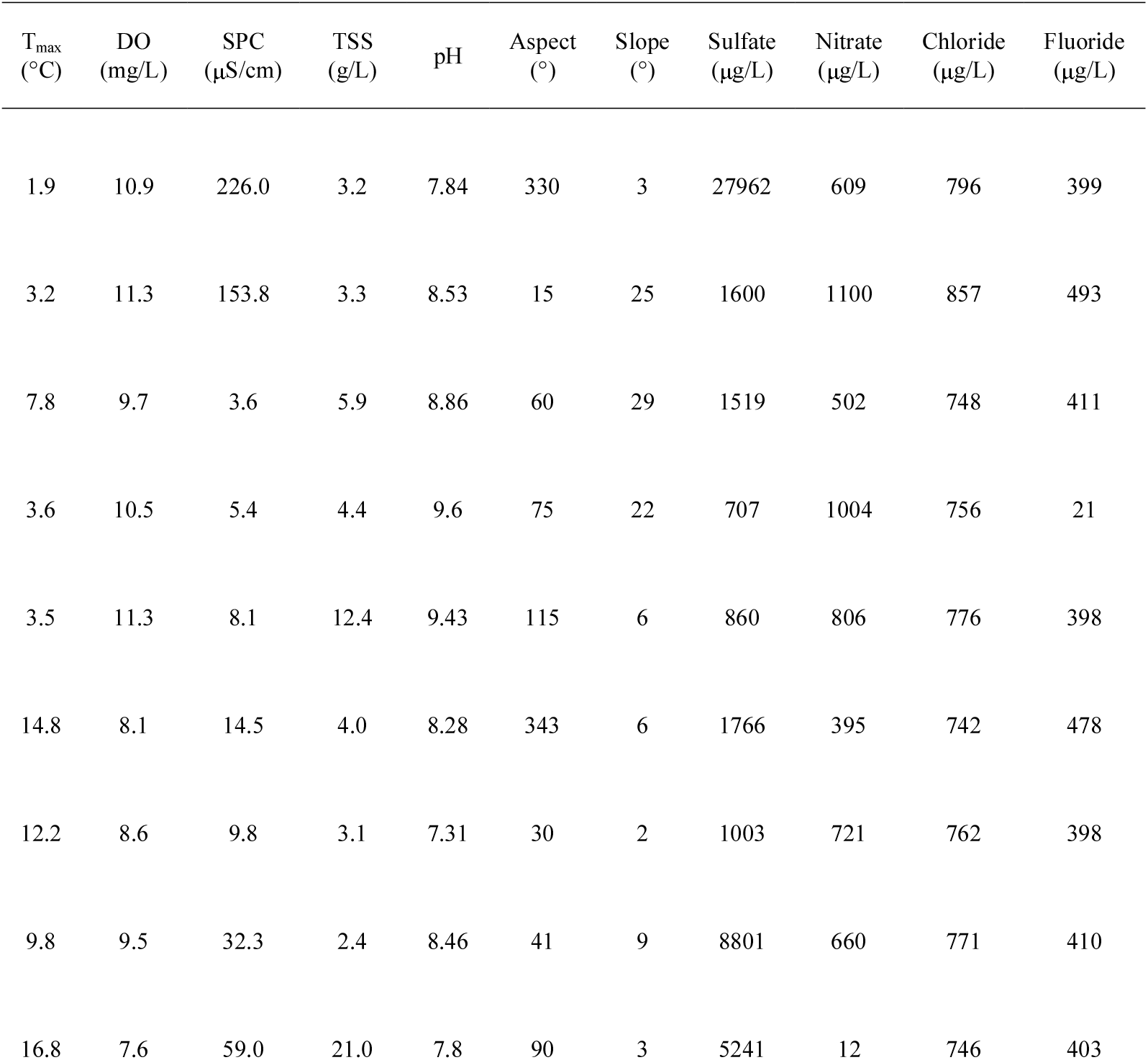
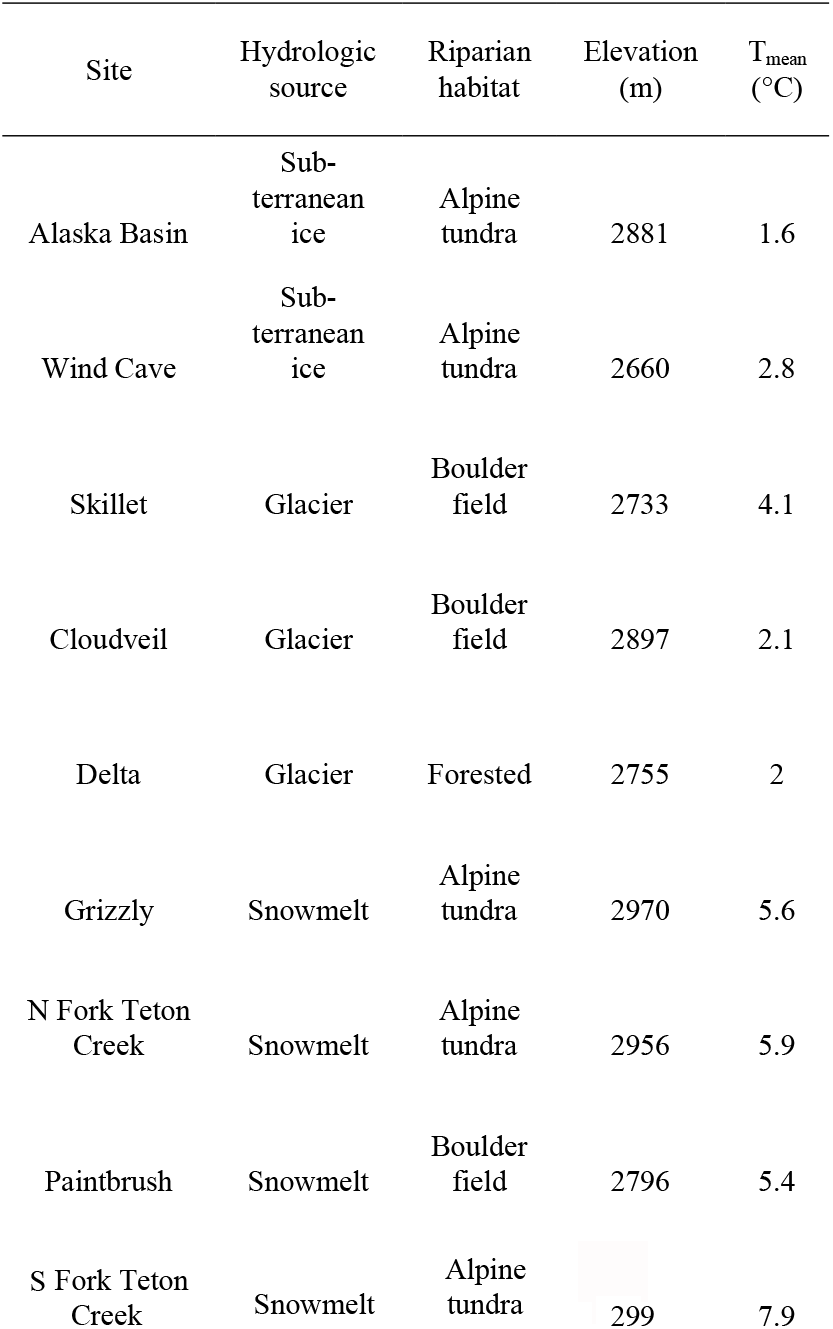
Site characteristics and environmental variables for alpine stream sites in the Teton Range, Wyoming.

**Figure 6.**
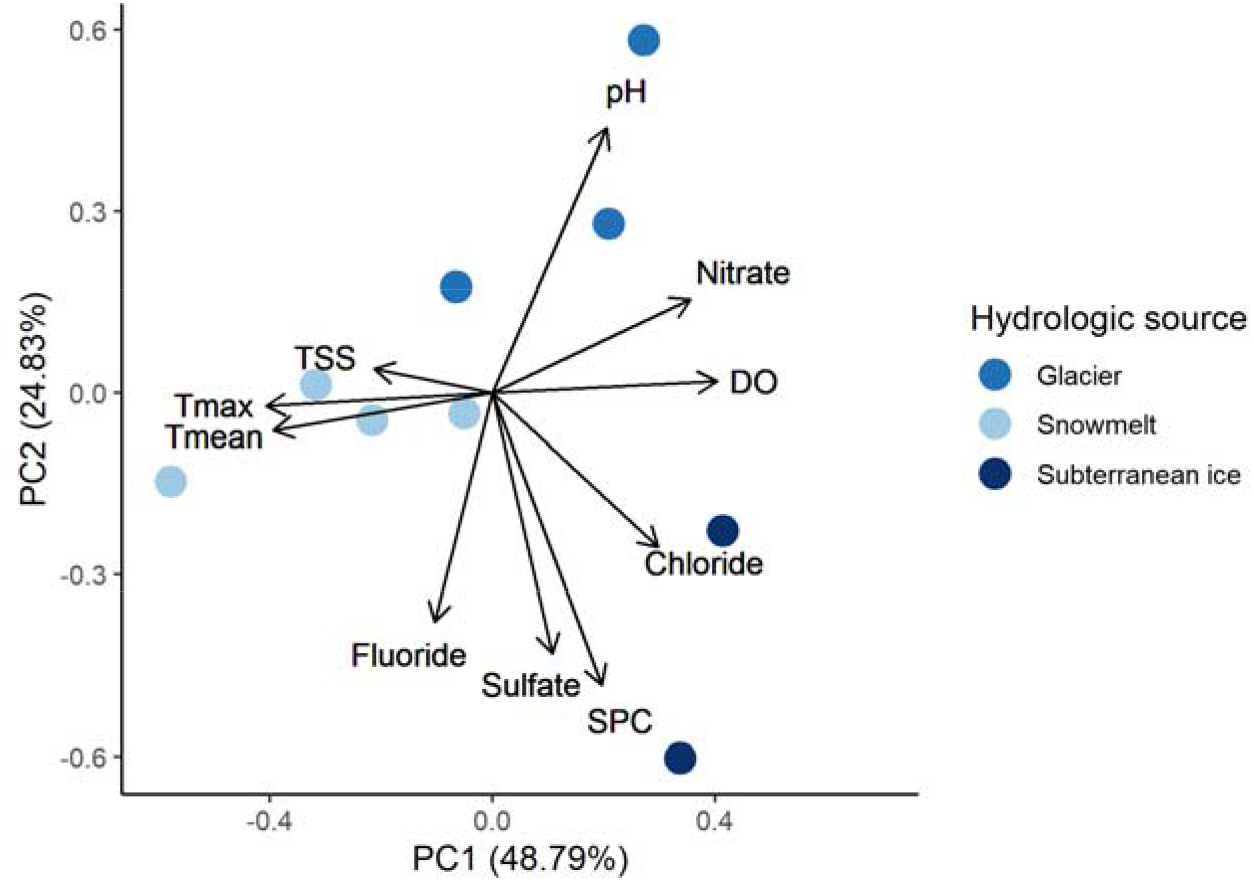
Principle component analyses showing environmental variation among sites with different hydrologic sources.

SPC (CPOM: -coefficient = 0.02, p-value = 0.002; *Hydrurus:* coefficient = 0.03, p-value < 0.001), T_mean_ (biofilm: coefficient = 0.47, p-value = 0.028; CPOM: -coefficient = 0.42, p-value = 0.044), chloride (*Hydrurus:* coefficient = 0.04, p-value = 0.004), and TSS (biofilm: coefficient = 32.43, p-value = 0.005; CPOM: coefficient = 33.35, p-value = 0.019) all had significant relationships with diet composition (Figure 7, Appendix 2 Table S2,). PC2 (biofilm: coefficient = 0.89, p-value = 0.012) also significantly influenced the diet composition and had a similar AICc value to individual environmental variables, while the model with hydrologic source had a much lower AICc value (Table 2). Combining environmental variables or environmental variables and hydrologic source increased the model fit, but the model did not converge for most combinations and sample size was too low to reliably combine explanatory variables. The biomass supported by each resource followed similar trends but with fewer significant relationships. Hydrologic source was still by far the best explanatory variable, with several individual variables that are known to be related to hydrologic source (Tronstad et al 2020) also having significant relationships (Figure 7), including SPC (CPOM: coefficient = 0.02, p-value = 0.010, *Hydrurus*: coefficient = 0.02, p-value < 0.001), TSS (biofilm: coefficient = 18.93, p-value = 0.034), and chloride (*Hydrurus*: coefficient = 0.04, p-value = 0.007). Examples of model checks are included in the appendix (Figures S11 and S12).

**Table 2.** Model selection using corrected Akaike information criterion (AICc). We observed a large increase in model fit between single environmental variables or the first PCA component (PC1) and models that included hydrologic source or multiple variables. Only models with significant effects are shown.

| Explanatory variables | AICc |
| --- | --- |
| Hydrologic source | -211.5 |
| SPC + Tmean | -200.7 |
| Hydrologic source + Tmax | -107.2 |
| SPC | 23.5 |
| Chloride | 29.4 |
| PC2 | 30.1 |
| TSS | 31.5 |
| Tmean | 31.7 |

**Figure 7.**
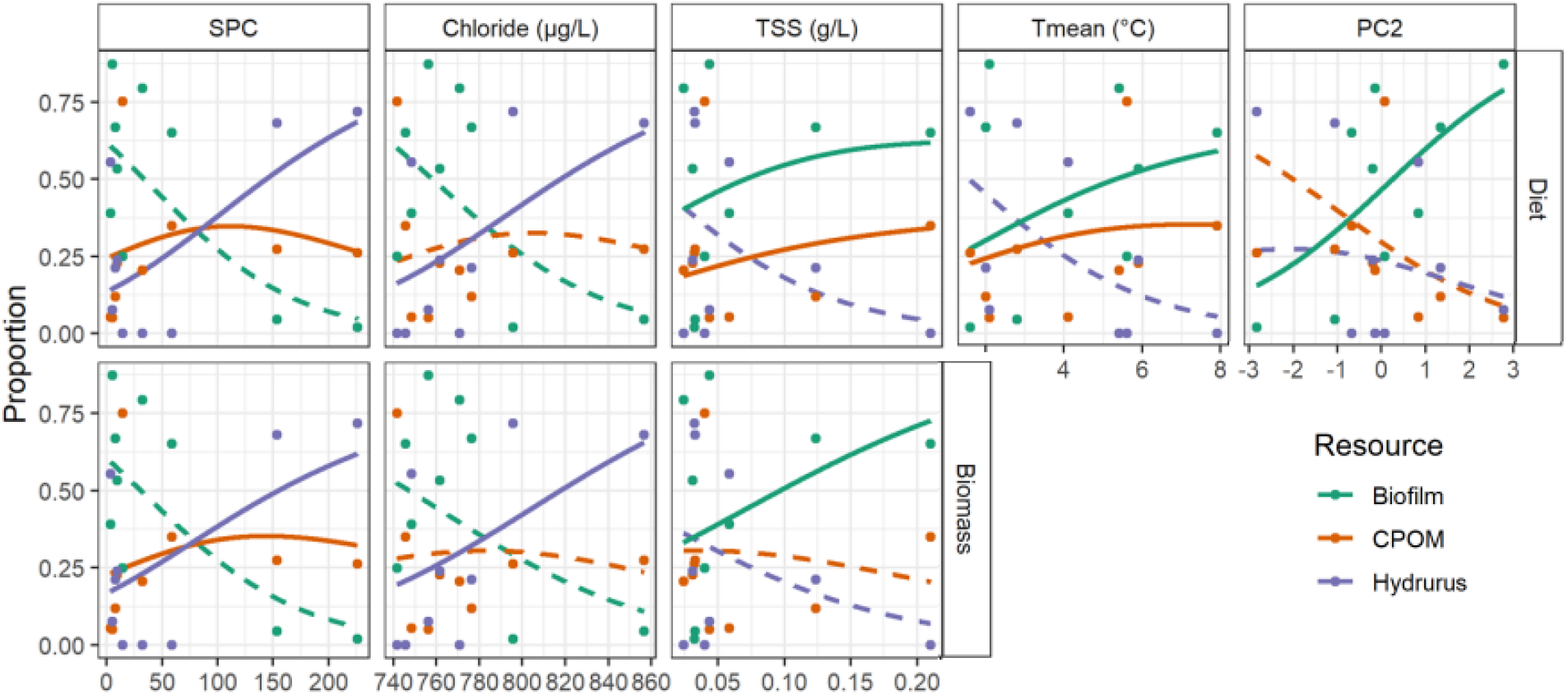
Models with significant relationships between diet or biomass proportion and environmental variables. Solid lines represent significant relationships, while dotted lines were not significant.

## Discussion

Understanding links between hydrologic source, imperiled taxa, and resource use in alpine streams strengthens our capacity to predict how climate change will impact these vulnerable ecosystems. In this study, we show that hydrologic source is a driver of food web structure, which expands on previous evidence about the important role of hydrology in shaping alpine diversity and community composition (Fell et al. 2017, Giersch et al. 2017). Overall, macroinvertebrate communities in alpine streams appear to rely heavily upon autochthonous resources, but large differences in the assimilation of biofilm and *Hydrurus* exist among hydrologic sources. This was particularly true for streams fed by subterranean ice—which are predicted to be most resilient to cryosphere recession (Brighenti et al. 2021)—as they exhibited the highest rates of *Hydrurus* assimilation. While we were not able to distinguish differences in resource use between snowmelt and glacier-fed streams with our sample size, previous research has shown that allochthonous diet contribution increases with decreasing environmental harshness in glacier-fed streams (Niedrist and Füreder 2018). Generally, hydrologic source explained diet compositions better than other environmental variables. Surprisingly, PCA components which summarized multiple environmental variables did not explain diet compositions better than individual variables.

Climate change is expected to influence the nutritional quality and availability of basal resources in alpine streams, which will in turn alter resource use in these communities. For instance, warmer water temperatures and decreased glacial influence are expected to increase the quantity of algal biofilms (Mosser & Brock 1976, Cauvy-Fraunié et al. 2016) and litter input is expected to increase as vegetation increases and shifts uphill (Xu et al. 2020, Emmet et al. 2019). However, nutrient limitation is an important consideration. Decreased glacier influence will likely decrease nitrogen availability, which could intensify nutrient limitation for autochthonous production (Slemmons et al. 2013). Additionally, litter decomposition (which increases nutritional quality) can also decrease with increasing stream intermittency resulting from reduced meltwater (Siebers et al. 2019) and nutrient limitation (Robinson & Gessner, 2000). The availability of *Hydrurus* may decrease in the future as the period when temperatures are too high for *Hydrurus* growth lengthens (Klaveness 2019, Hieber et al. 2001). As temperatures rise, currently abundant algae could be outcompeted by filamentous algae with lower nutritional quality (Oleksy et al. 2021, Brett et al. 1997). Although we lack a complete understanding of how climate change will influence food resources, there is potential for the quality of allochthonous resources, and the quantity and quality of autochthonous resources for stream macroinvertebrates to be reduced.

### *Trophic flexibility and the role of* Hydrurus *in alpine food webs*

Trophic flexibility can increase the resilience of organisms and communities to environmental change (Saint-Béat et al. 2015, Bartley et al. 2019). We observed a prevalence of trophic flexibility in alpine streams across hydrologic sources, with individual taxa and communities capitalizing on different resources among sites and seemingly unconstrained by feeding morphology. The low within-stream variability in resource use demonstrates that invertebrate communities in the Teton Range are consuming the same food resources instead of specializing, suggesting low pressure from competition (Mihuc 1997). Thus, it appears that at present, competition for resources is not an important driver of macroinvertebrate distributions in Teton alpine streams. Furthermore, community diet variance at a given site did not increase with taxonomic diversity, indicating that whole communities are targeting the same resources, even in more diverse snowmelt-fed streams. Trophic flexibility likely helps enable many invertebrate taxa to maintain populations in diverse alpine stream habitats, as has been seen in other ecosystems with large environmental gradients (Schalk et al. 2017, Leclerc et al. 2021).

Gut content analysis allowed us to observe flexible feeding at a smaller scale, including predatory behavior in taxa that are usually primary consumers (e.g., black fly larvae, Simuliidae), potentially due to low quality resources (Diehl 2003). We also observed consumption of algae by predators, which may result from low prey availability (Coll & Guershon 2002) and could reduce predation stress on primary consumers. Trophic flexibility by macroinvertebrate communities in alpine streams suggests that these taxa do not conform to traditional functional feeding groups and that population growth may be limited more by the harsh environment than by food availability.

Despite the benefits of flexible feeding strategies, they may not be sufficient to stabilize alpine food webs if the degradation of a key resource such as *Hydrurus* occurs. *Hydrurus* is an especially valuable resource because of its high fatty acid content (Klaveness 2017) and abundance early in the growing season (Rott et al. 2006). Alterations in the phenology of food resources due to climate change will have detrimental effects on many consumers worldwide (Parmesan 2006). If the growth of *Hydrurus* is limited by stream temperatures rising in the spring (Klaveness 2019, Hieber et al. 2001), taxa may struggle to acquire the nutrients needed early in the growing season. Thus, even if overall primary productivity increases in alpine streams, the decreased growth of an alga that is abundant before snow and ice recedes may limit macroinvertebrate development. We also observed lower consumption of *Hydrurus* in warmer snowmelt-fed streams when *Hydrurus* may be senescing, suggesting that it may be a lower quality resource when dying back. The typically distinct carbon isotope signature of *Hydrurus* provides a valuable opportunity to observe changes in its importance over time or to detect its food web signature when streams are inaccessible in the spring and *Hydrurus* is most abundant.

Because *Hydrurus* appears to be an integral part of alpine stream food webs and changes in its availability will likely have larger ramifications than other food resources, its presence and persistence could be relevant for identifying refugia where meltwater biodiversity may persist under climate change. Rock glaciers, the most common type of subterranean ice, are ∼10 times more abundant than traditional surface glaciers in the contiguous US (Johnson 2018). Additionally, rock glaciers are predicted to melt slower than glaciers because of the insulation provided by layers of rock and debris (Anderson et al. 2018, Brighenti et al. 2021). Thus, our finding that macroinvertebrate communities in these streams had the largest assimilation of *Hydrurus* in August suggests that it may persist longer into the growing season compared to streams with glacier or snowmelt sources. Subterranean ice-fed streams lack the high turbidity and scouring of glacier-fed streams which limits algal productivity during the summer (Rott et al. 2006, Hieber et al. 2001, Uehlinger et al. 2009), while still maintaining relatively high nitrogen (Fegel et al. 2016) and the cold temperatures necessary for *Hydrurus* (Klaveness 2017).

*Hydrurus* may therefore be more stable in subterranean ice-fed streams than those fed by other hydrologic sources. Given that subterranean ice-fed streams in the Teton Range maintain diverse macroinvertebrate assemblages relative to other stream types, including rare cold-water taxa (e.g., *L. tetonica* and *Zapada glacier*, Tronstad et al. 2020), the stability of *Hydrurus* in these streams may contribute to their long-term potential as high mountain refugia for aquatic biodiversity. This combination of subterranean ice meltwater and abundance of high-quality resources may also explain why cold-water biodiversity has been observed persisting in some reaches of other montane regions long after glaciers have receded (Muhlfeld et al. 2020).

### Implications for vulnerable taxa in the Teton Range

In the Teton Range, two stoneflies—*L. tetonica* and *Z. glacier*—are of conservation and management interest as they are either ESA-listed (*Z. glacier*) or the close congeneric of another listed species (*L. tetonica*). However, both species are poorly studied with basic aspects of their life history largely unknown, including their dietary needs, which makes management planning difficult. We show that both taxa can switch from biofilm to *Hydrurus* dominated diets, and *Zapada* consumed more allochthonous resources than *L. tetonica*. These taxa were also the only taxa observed to consume wood at any of our sites. In one glacier-fed stream, our analyses indicated that *L. tetonica* may exhibit predatory behavior. This finding aligns with a recent study of its congener—*L. tumana—*which can exhibit cannibalism in captivity (Shah et al. 2022). These taxa are most abundant in the coldest sections of alpine streams, making shifts to higher elevations unlikely (Giersch et al. 2017). The high consumption of *Hydrurus* by *L. tetonica, Zapada*, and co-occurring species in our study, including in streams fed by subterranean ice, further supports the idea that these streams may act as climate refugia (Brighenti et al. 2021). By pairing prior knowledge of species distributions with their trophic ecology and predictions about which hydrologic sources will be most resilient to climate change, managers are left with a clearer picture of what habitats warrant protection for limiting the magnitude of biodiversity loss as the cryosphere fades.

### Conclusion

The hydrology of alpine ecosystems is changing rapidly, and the communities within streams fed by distinct hydrologic sources will likely be impacted by these changes at different rates. Flexible feeding strategies may allow the persistence of alpine taxa until other impacts of climate change, including invasive species or stream intermittency, become primary limiting factors. In addition to many other factors that make them suitable as potential climate refugia, subterranean ice features also support abundant *Hydrurus* populations, a key food resource for many taxa including vulnerable species. Further study is necessary to test these results in additional mountain ranges and geographic regions with different seasonal resource dynamics, and over broader spatial scales. This study advances our understanding of how alpine stream ecosystems function across different hydrologic sources and provides baseline data for further exploring what the future may hold for these imperiled communities.

## Acknowledgements

This project was supported with funding from the Meg and Bert Raynes Wildlife Fund, the University of Wyoming-NPS Small Grants Program, the Vern Bressler Fisheries Fund, the University of Wyoming, the Jackson Hole One Fly Scholarship, and the National Science Foundation (OIA-2019596). Taylor Price, Shannon Weld, Isabella Errigo, Anna Eichert, Mikayla Castillon, Macy Jacobson, Casey Brucker, Dana Hasert and Maddison Kopsa assisted with field collections. Additional lab help came from Katie Bearden, Angela Zhu, and Jaide Phelps, with special thanks to Chandelle MacDonald and others at the University of Wyoming Stable Isotope Facility. The Collins laboratory group and Daniel Laughlin provided important feedback on the manuscript.

