## Supplementary Materials for "Trophic flexibility and hydrology structure alpine stream food webs: Implications for a fading cryosphere"

**Appendix 1**

*Trophic enrichment factor selection*

One primary challenge for using stable isotope analysis to analyze food webs is selecting an appropriate trophic enrichment factor (TEF). An appropriate nitrogen TEF is important in alpine streams to correctly allocate diet proportions, especially between CPOM and *Hydrurus*. Uncertainty in the TEF may have resulted in an overestimate of the diet contribution of CPOM in this study. Empirically determining TEFs for alpine systems would be logistically challenging, so we relied on estimates from meta-analyses. In our isotope mixing models we used a nitrogen TEF estimated by Bunn et al. (2013) specific for invertebrate herbivores in streams and rivers that has been used in previously in alpine stream studies (Niedrist & Füreder 2018). The estimate from the Bunn (2013) paper is lower than the general estimate from Post (2002) that is commonly used in isotope studies and is also lower than estimates including predators. While an estimate for invertebrate herbivores is more appropriate than a general estimate (Brauns et al. 2018, Caut et al. 2009, McCutchan et al. 2003) or one that includes higher trophic levels (Bunn et al. 2013, Jardine et al. 2005), continued investigation into TEFs specifically targeted toward taxonomic groups and different ecosystems.

**Appendix 2**

Table S1. Functional feeding groups for the taxa included in the isotope food web analysis. FFGs were compiled from Merritt et al. (2008).

| **Family** | **Genus** | **Functional feeding group** |
| --- | --- | --- |
| Ameletidae |  | Scrapers; collectors-gatherers |
| Apatainidae | Allomyia | Shredders-herbivores; scrapers |
| Baetidae | Heterocloeon | Scrapers |
|  | Baetis | Collector-gatherers; facultative scrapers |
| Blephraceridae | Agathon | Scrapers |
| Chloroperlidae | Sweltsa | Predators; facultative collectors-gatherers |
| Empedidae | Clinocera | Unknown |
| Ephemerellidae | Drunella | Scrapers; facultative predators |
| Heptageniidae | Rithrogena | Scrapers; facultative collectors-gatherers |
|  | Epeorus | Scrapers; facultative collectors-gatherers |
|  | Cinygmula | Scrappers; facultative collectors-gatherers |
| Limnephilidae | Homphylax | Shredders-detritivores |
| Nemouridae |  | Shredders-detritivores; facultative collectors-gatherers |
|  | Lednia | Unknown |
|  | Zapada | Shredders-detritivores |
| Perlodidae | Megarcys | Predators |
| Rhyacophilidae | Rhyacophila | Predators; a few scrapers, collectors-gatherers; shredders-herbivores |
| Simullidae |  | Collectors-filterers; facultative scrapers and predators |
| Tipulidae | Tipula | Shredders-detritivores; facultative shredders-herbivores, and collectors-gatherers |
| Non-Tanypodinae midges |  | Collectors-gatherers and filterers |

Table S2. Model estimates and 95% confidence intervals for diet proportion models. Significant (α = 0.05) responses are indicated with an asterisk.

| **Variables** | **Resource** | **Estimate** | **2.5%** | **97.5%** | **P-value** |  |
| --- | --- | --- | --- | --- | --- | --- |
| Specific conductivity | Biofilm | 0.01 | -0.01 | 0.02 | 0.280 |  |
|  | CPOM | 0.02 | 0.01 | 0.03 | 0.002 | * |
|  | *Hydrurus* | 0.03 | 0.01 | 0.04 | 0.000 | * |
| Temperature (mean) | Biofilm | 0.47 | 0.05 | 0.89 | 0.028 | * |
|  | CPOM | 0.42 | 0.01 | 0.83 | 0.044 | * |
|  | *Hydrurus* | 0.00 | -0.36 | 0.37 | 0.986 |  |
| Temperature (max) | Biofilm | 0.13 | 0.00 | 0.26 | 0.051 |  |
|  | CPOM | 0.15 | 0.00 | 0.30 | 0.052 |  |
|  | *Hydrurus* | -0.04 | -0.17 | 0.10 | 0.600 |  |
| pH | Biofilm | 1.70 | -0.06 | 3.46 | 0.059 |  |
|  | CPOM | 0.41 | -0.83 | 1.64 | 0.519 |  |
|  | *Hydrurus* | 0.87 | -0.43 | 2.17 | 0.189 |  |
| Nitrate | Biofilm | 0.00 | 0.00 | 0.00 | 0.429 |  |
|  | CPOM | 0.00 | 0.00 | 0.00 | 0.337 |  |
|  | *Hydrurus* | 0.00 | 0.00 | 0.00 | 0.665 |  |
| Total suspended solids | Biofilm | 32.43 | 9.75 | 55.10 | 0.005 | * |
|  | CPOM | 33.35 | 5.58 | 61.13 | 0.019 | * |
|  | *Hydrurus* | 17.74 | -2.77 | 38.26 | 0.090 |  |
| Elevation | Biofilm | 0.00 | -0.01 | 0.01 | 0.627 |  |
|  | CPOM | 0.00 | -0.01 | 0.01 | 0.583 |  |
|  | *Hydrurus* | 0.00 | -0.01 | 0.00 | 0.296 |  |
| Dissolved oxygen | Biofilm | -0.48 | -1.02 | 0.06 | 0.082 |  |
|  | CPOM | -0.55 | -1.19 | 0.10 | 0.097 |  |
|  | *Hydrurus* | 0.16 | -0.40 | 0.71 | 0.581 |  |
| Slope | Biofilm | 0.00 | -0.07 | 0.07 | 0.921 |  |
|  | CPOM | -0.02 | -0.09 | 0.05 | 0.642 |  |
|  | *Hydrurus* | 0.05 | -0.04 | 0.13 | 0.294 |  |
| Aspect | Biofilm | 0.00 | -0.01 | 0.00 | 0.349 |  |
|  | CPOM | 0.00 | -0.01 | 0.01 | 0.593 |  |
|  | *Hydrurus* | 0.00 | -0.01 | 0.01 | 0.779 |  |
| Chloride | Biofilm | 0.01 | -0.02 | 0.04 | 0.541 |  |
|  | CPOM | 0.03 | 0.00 | 0.06 | 0.054 |  |
|  | *Hydrurus* | 0.04 | 0.01 | 0.07 | 0.004 | * |
| Sulfate | Biofilm | 0.00 | 0.00 | 0.00 | 0.310 |  |
|  | CPOM | 0.00 | 0.00 | 0.00 | 0.108 |  |
|  | *Hydrurus* | 0.00 | 0.00 | 0.00 | 0.055 |  |
| Principal component 1 | Biofilm | -0.35 | -0.71 | 0.01 | 0.058 |  |
|  | CPOM | -0.35 | -0.74 | 0.05 | 0.083 |  |
|  | *Hydrurus* | 0.08 | -0.30 | 0.45 | 0.691 |  |
| Principal component 2 | Biofilm | 0.89 | 0.19 | 1.60 | 0.012 | * |
|  | CPOM | 0.27 | -0.33 | 0.87 | 0.371 |  |
|  | Hydrurus | 0.46 | -0.26 | 1.18 | 0.210 |  |
| Hydrologic source |  |  |  |  |  |  |
| Glacier:Snowmelt | Biofilm | 0.39 | -1.19 | 1.97 | 0.630 |  |
|  | CPOM | -0.76 | -2.31 | 0.80 | 0.340 |  |
|  | *Hydrurus* | 1.18 | -0.36 | 2.72 | 0.130 |  |
| Glacier:Subterranean ice | Biofilm | -2.02 | -3.84 | -0.19 | 0.030 | * |
|  | CPOM | -4.94 | -6.75 | -3.14 | < 0.001 | * |
|  | *Hydrurus* | -5.15 | -6.97 | -3.33 | < 0.001 | * |
| Snowmelt:Subterranean ice | Biofilm | -2.41 | -4.14 | -0.67 | 0.007 | * |
|  | CPOM | -4.18 | -5.92 | -2.45 | < 0.001 | * |
|  | *Hydrurus* | -5.15 | -6.97 | -3.33 | < 0.001 | * |
| Hydrologic source + temperature (max) |  |  |  |  |  |  |
| Glacier:Subterranean ice | Biofilm | 0.86 | -2.02 | 3.75 | 0.557 |  |
|  | CPOM | -2.73 | -5.46 | 0.01 | 0.050 |  |
|  | *Hydrurus* | -3.08 | -5.55 | -0.61 | 0.015 | * |
| Snowmelt:Subterranean ice | Biofilm | 3.85 | -1.97 | 9.67 | 0.194 |  |
|  | CPOM | -0.31 | -5.22 | 4.60 | 0.900 |  |
|  | *Hydrurus* | -2.34 | -6.79 | 2.12 | 0.305 |  |
| Temperature (max) | Biofilm | -0.45 | -0.90 | -0.01 | 0.045 | * |
|  | CPOM | -0.24 | -0.62 | 0.14 | 0.210 |  |
|  | *Hydrurus* | -0.25 | -0.63 | 0.12 | 0.182 |  |
| Specific conductivity + temperature (mean) |  |  |  |  |  |  |
| Specific conductivity | Biofilm | 0.00 | -0.01 | 0.01 | 0.830 |  |
|  | CPOM | 0.02 | 0.01 | 0.03 | 0.004 | * |
|  | *Hydrurus* | 0.02 | 0.01 | 0.03 | 0.001 | * |
| Temperature | Biofilm | -0.70 | -1.40 | 0.01 | 0.052 |  |
|  | CPOM | -0.38 | -0.92 | 0.16 | 0.172 |  |
|  | *Hydrurus* | -0.58 | -0.97 | -0.19 | 0.004 | * |

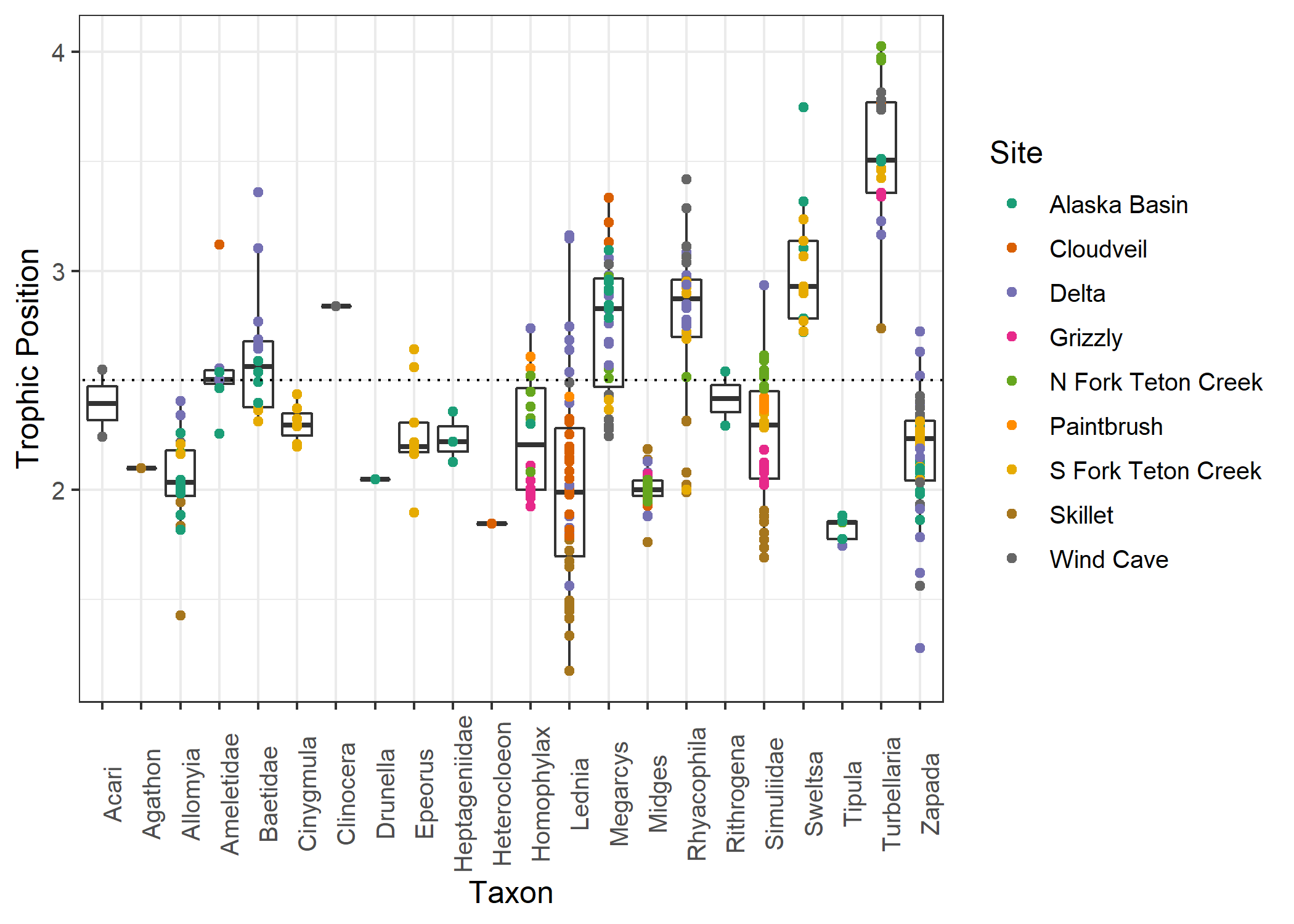

Figure S1. Trophic position estimates for each individual across all sites calculated using a one-baseline formula. Either *Allomyia* or midges were used as the baseline, and we used a TEF of 3.4 (Post 2002). The dotted line represents a trophic position of 2.5, which we used as a cutoff for including individuals in our isotope mixing models as primary consumers.

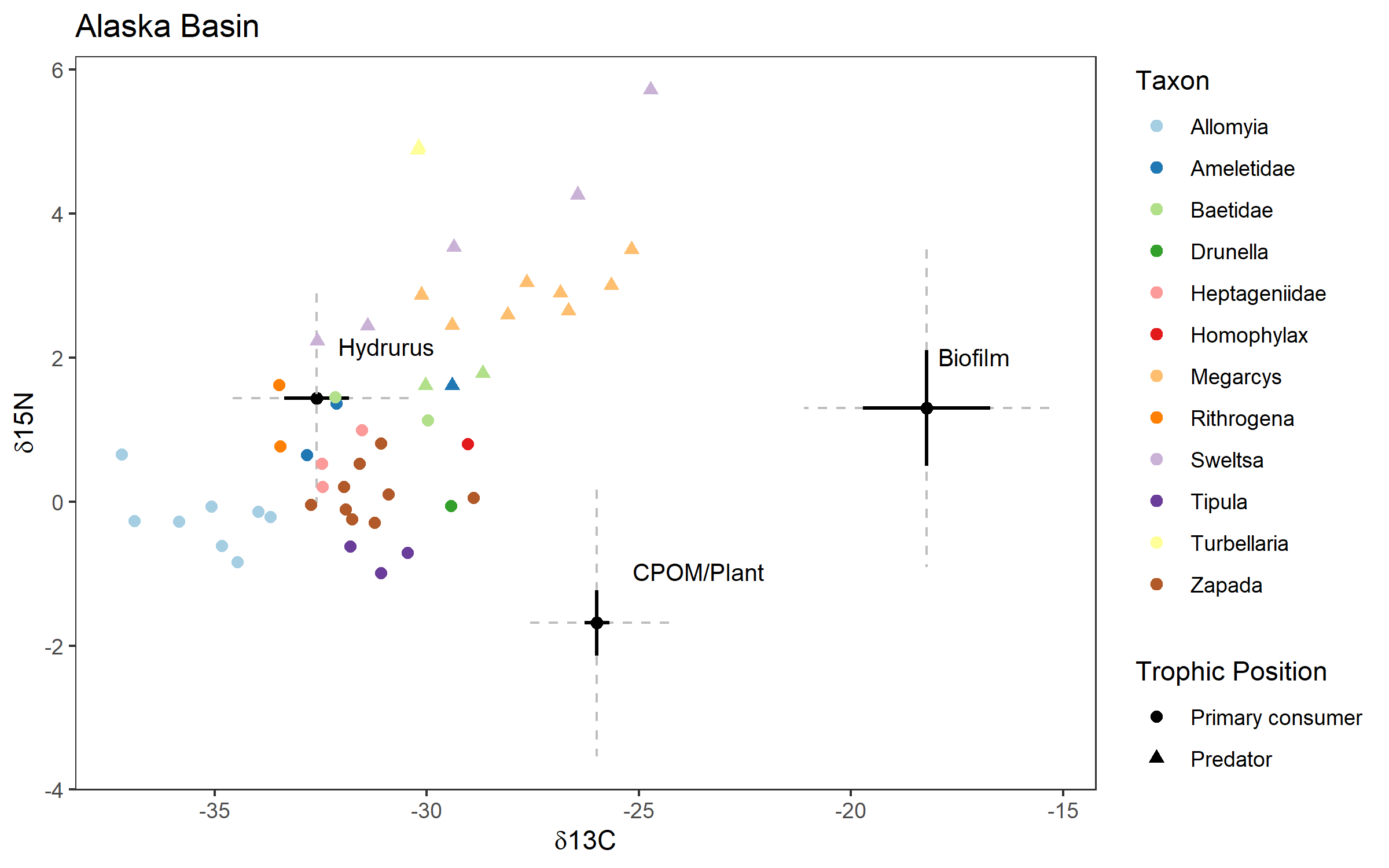

Figure S2. Raw isotope values for Alaska Basin (subterranean ice-fed). Resources are represented as means and standard errors and individual invertebrates are represented as points. Resources were shifted by the TEF (0.4 ‰ ± 1.4 for δ^13^C and 1.4 ± 1.4 for δ^15^N), and grey dashed lines indicate the additional error added by the TEFs.

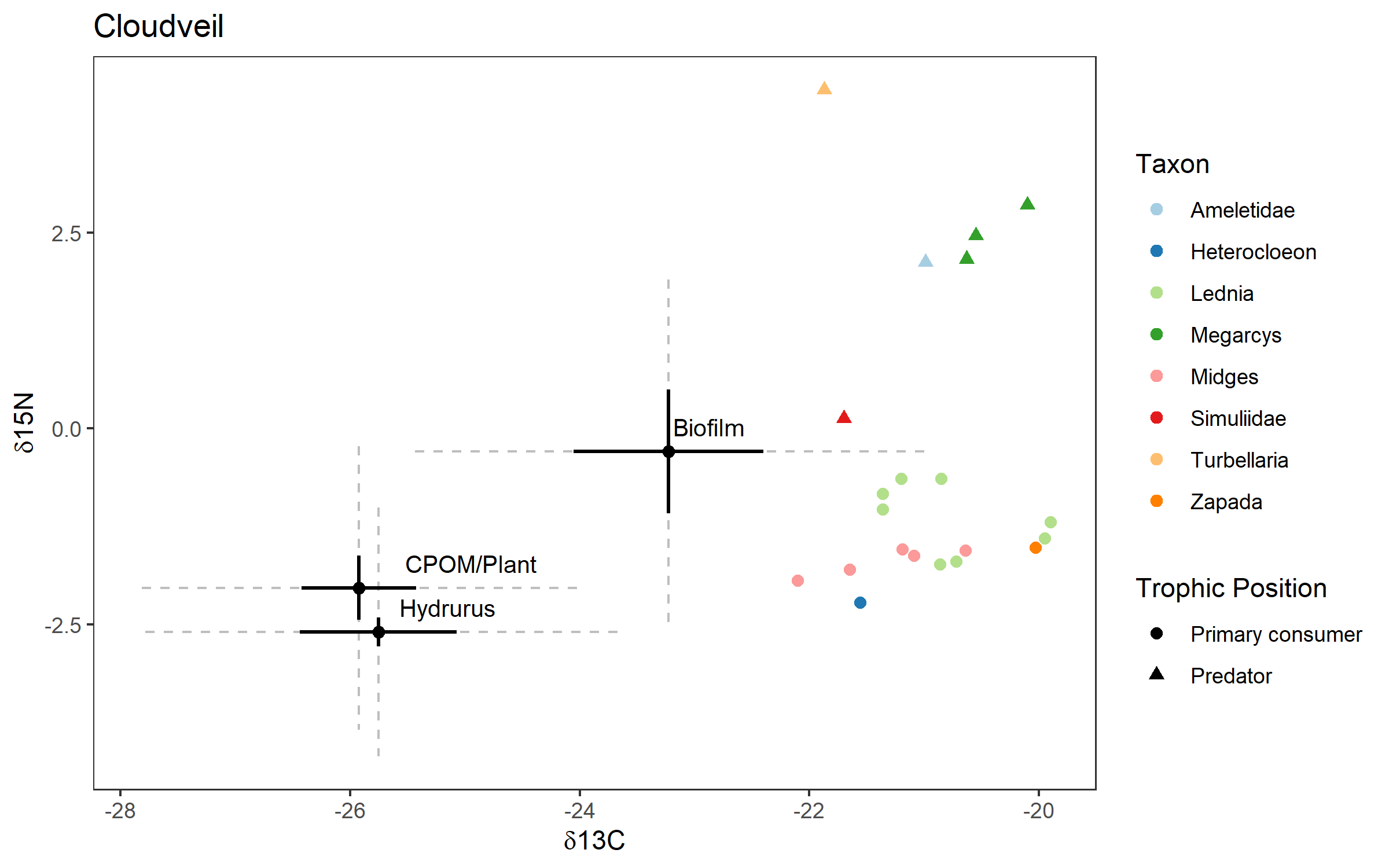

Figure S3. Raw isotope values for Cloudveil (glacier-fed). Resources are represented as means and standard errors and individual invertebrates are represented as points. Resources were shifted by the TEF (0.4 ‰ ± 1.4 for δ^13^C and 1.4 ± 1.4 for δ^15^N), and grey dashed lines indicate the additional error added by the TEFs.

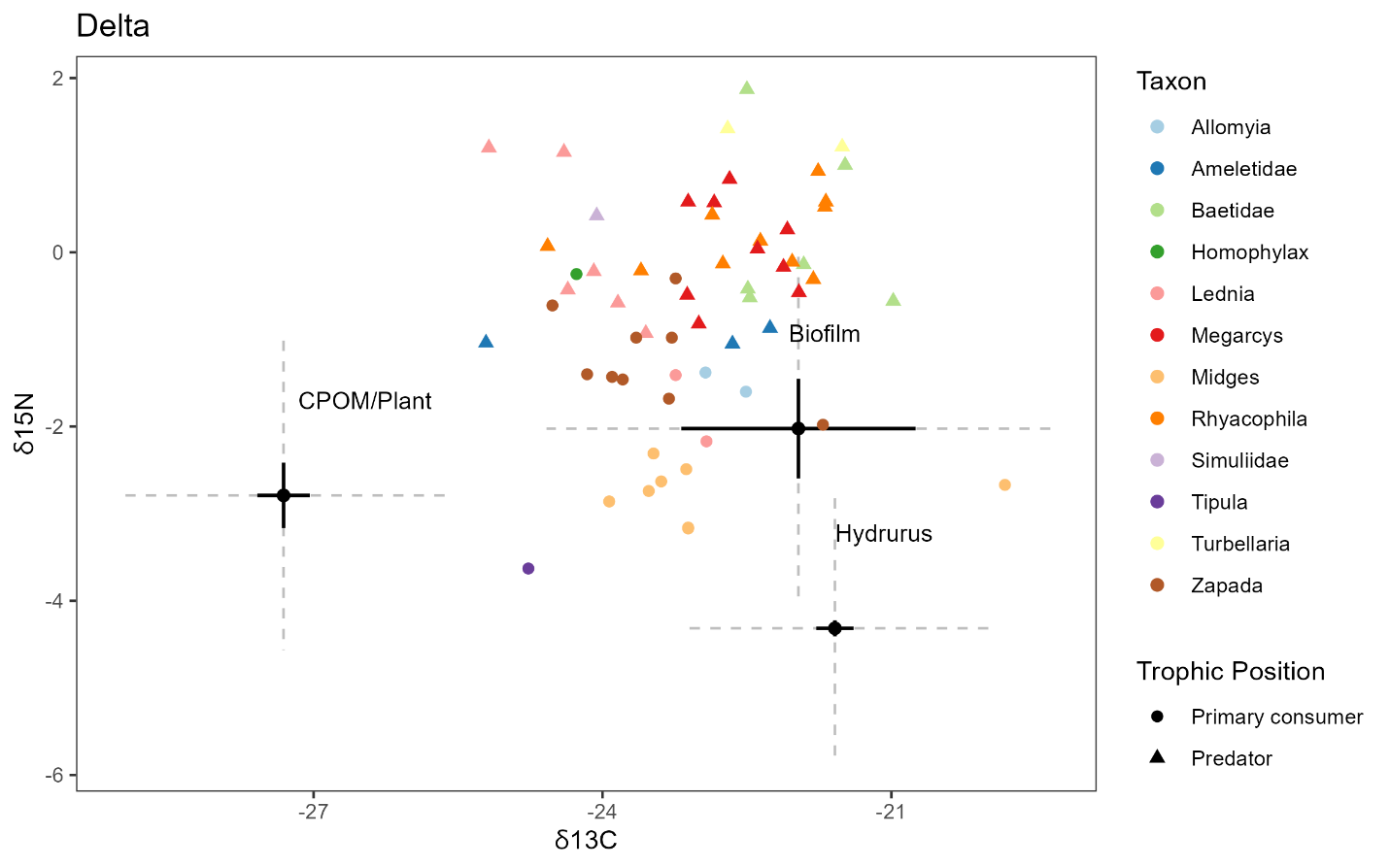

Figure S4. Raw isotope values for Delta (glacier-fed). Resources are represented as means and standard errors and individual invertebrates are represented as points. Resources were shifted by the TEF (0.4 ‰ ± 1.4 for δ^13^C and 1.4 ± 1.4 for δ^15^N), and grey dashed lines indicate the additional error added by the TEFs.

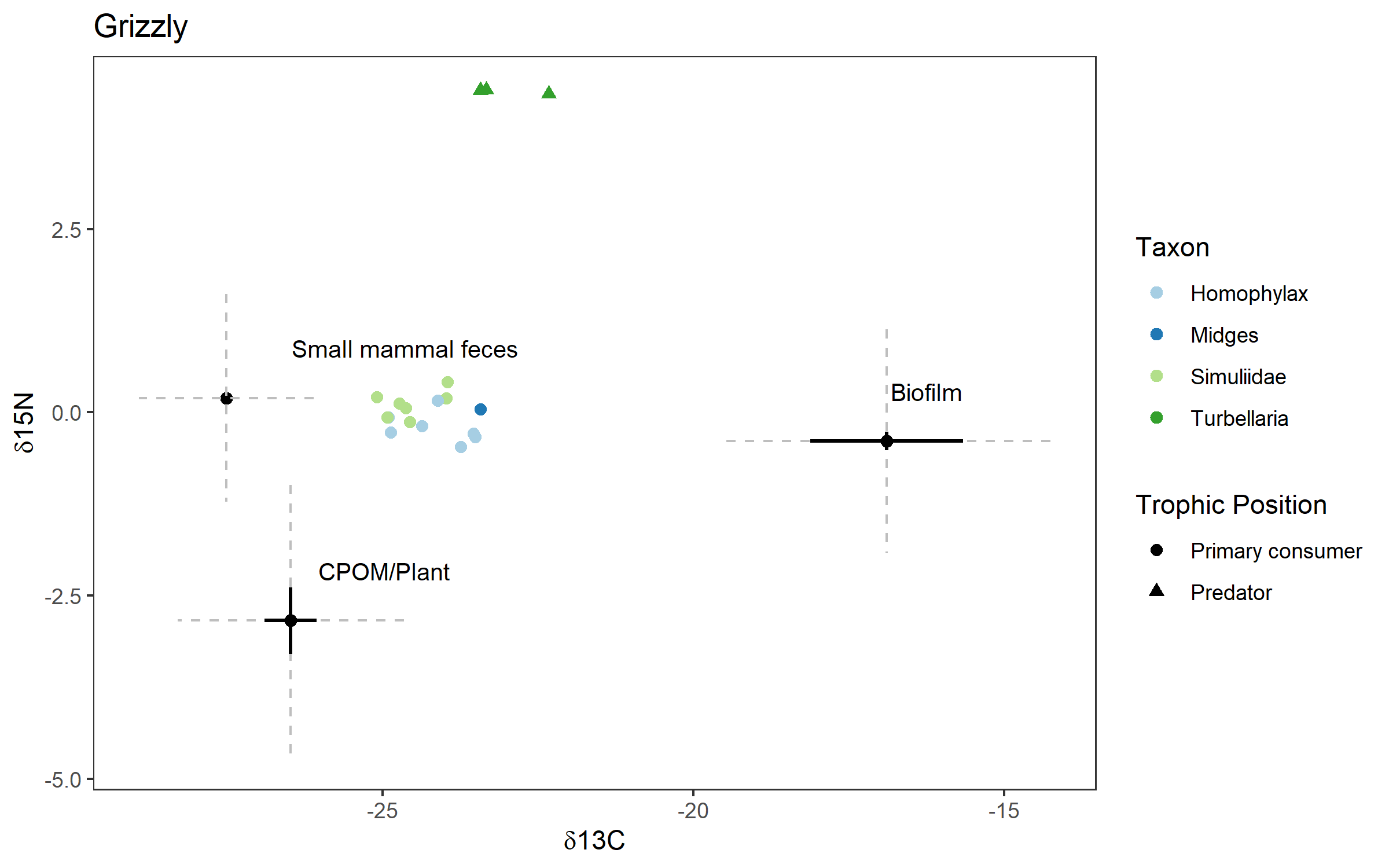

Figure S5. Raw isotope values for Grizzly (snowmelt-fed). Resources are represented as means and standard errors and individual invertebrates are represented as points. Resources were shifted by the TEF (0.4 ‰ ± 1.4 for δ^13^C and 1.4 ± 1.4 for δ^15^N), and grey dashed lines indicate the additional error added by the TEFs.

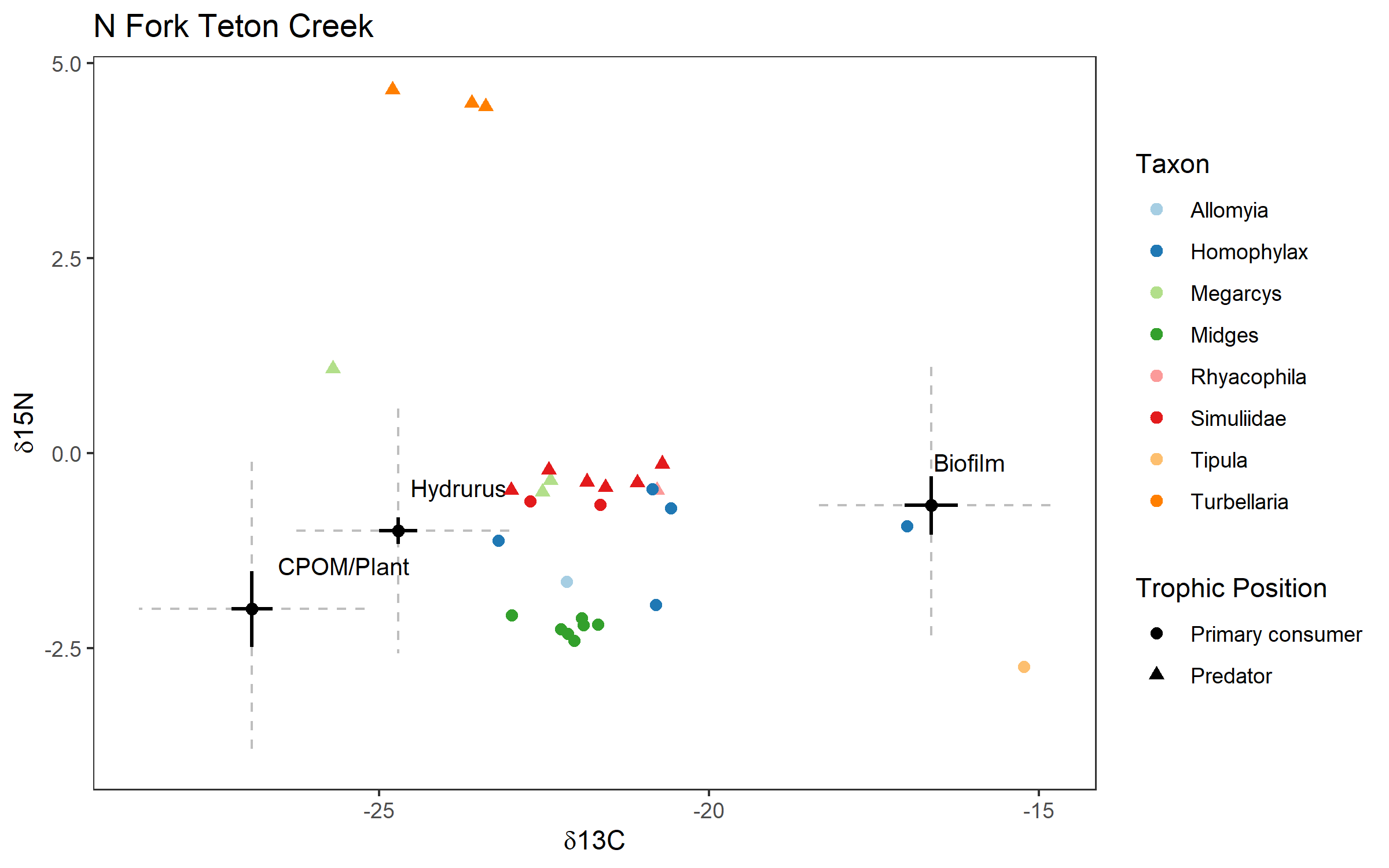

Figure S6. Raw isotope values for N Fork Teton Creek (snowmelt-fed). Resources are represented as means and standard errors and individual invertebrates are represented as points. Resources were shifted by the TEF (0.4 ‰ ± 1.4 for δ^13^C and 1.4 ± 1.4 for δ^15^N), and grey dashed lines indicate the additional error added by the TEFs.

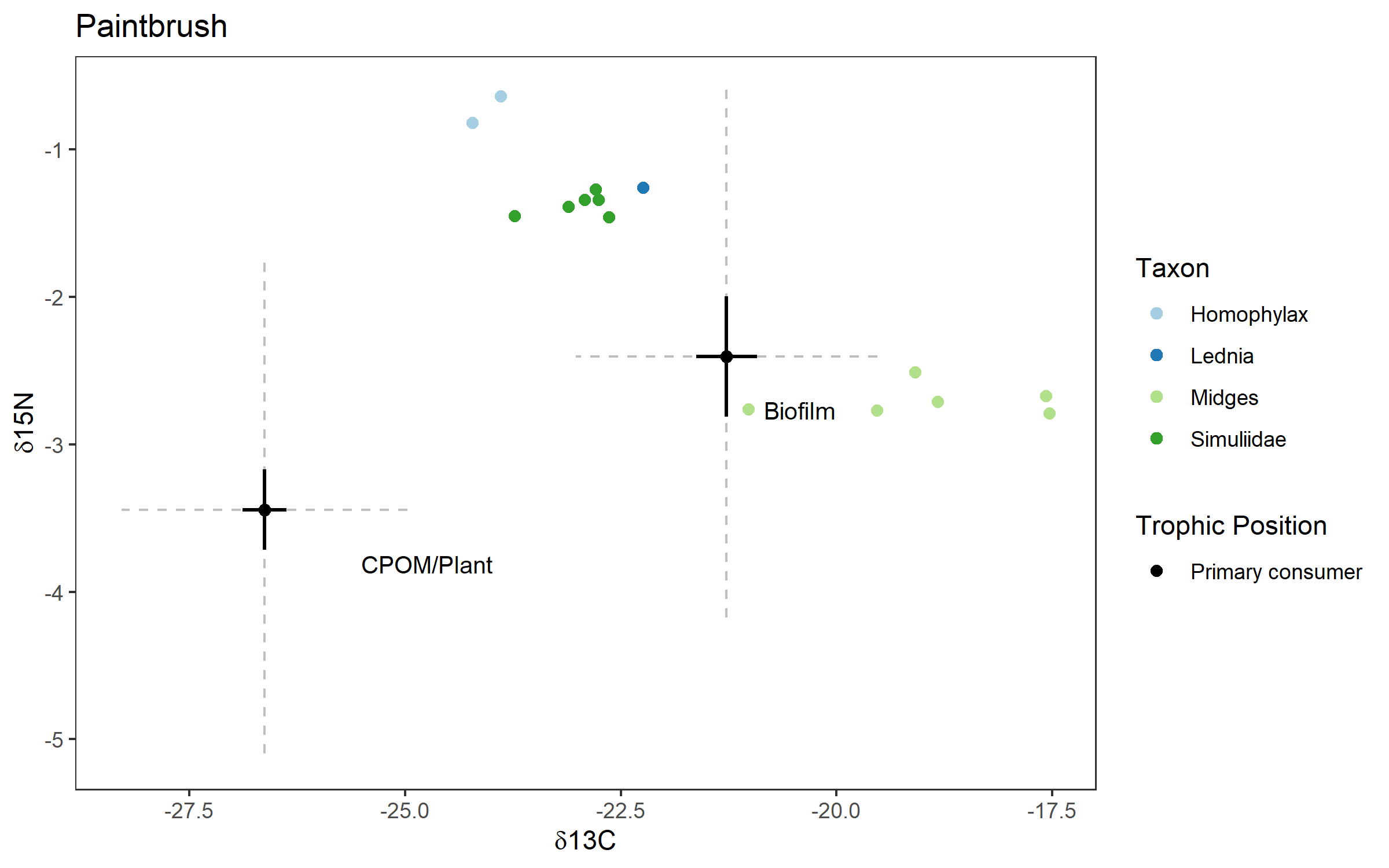

Figure S7. Raw isotope values for Paintbrush (snowmelt-fed). Resources are represented as means and standard errors and individual invertebrates are represented as points. Resources were shifted by the TEF (0.4 ‰ ± 1.4 for δ^13^C and 1.4 ± 1.4 for δ^15^N), and grey dashed lines indicate the additional error added by the TEFs.

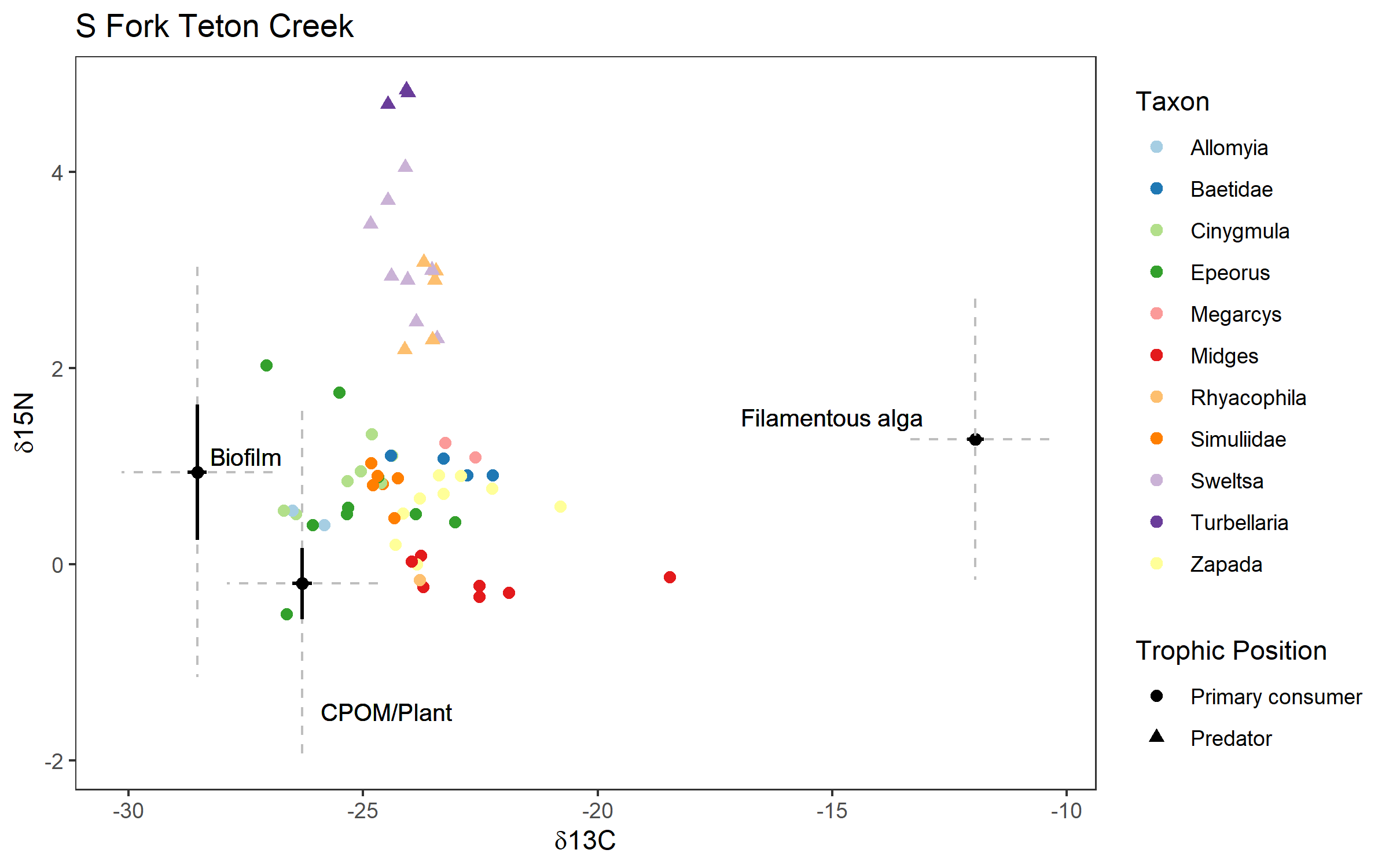

Figure S8. Raw isotope values for South Fork Teton Creek (snowmelt-fed). Resources are represented as means and standard errors and individual invertebrates are represented as points. Resources were shifted by the TEF (0.4 ‰ ± 1.4 for δ^13^C and 1.4 ± 1.4 for δ^15^N), and grey dashed lines indicate the additional error added by the TEFs.

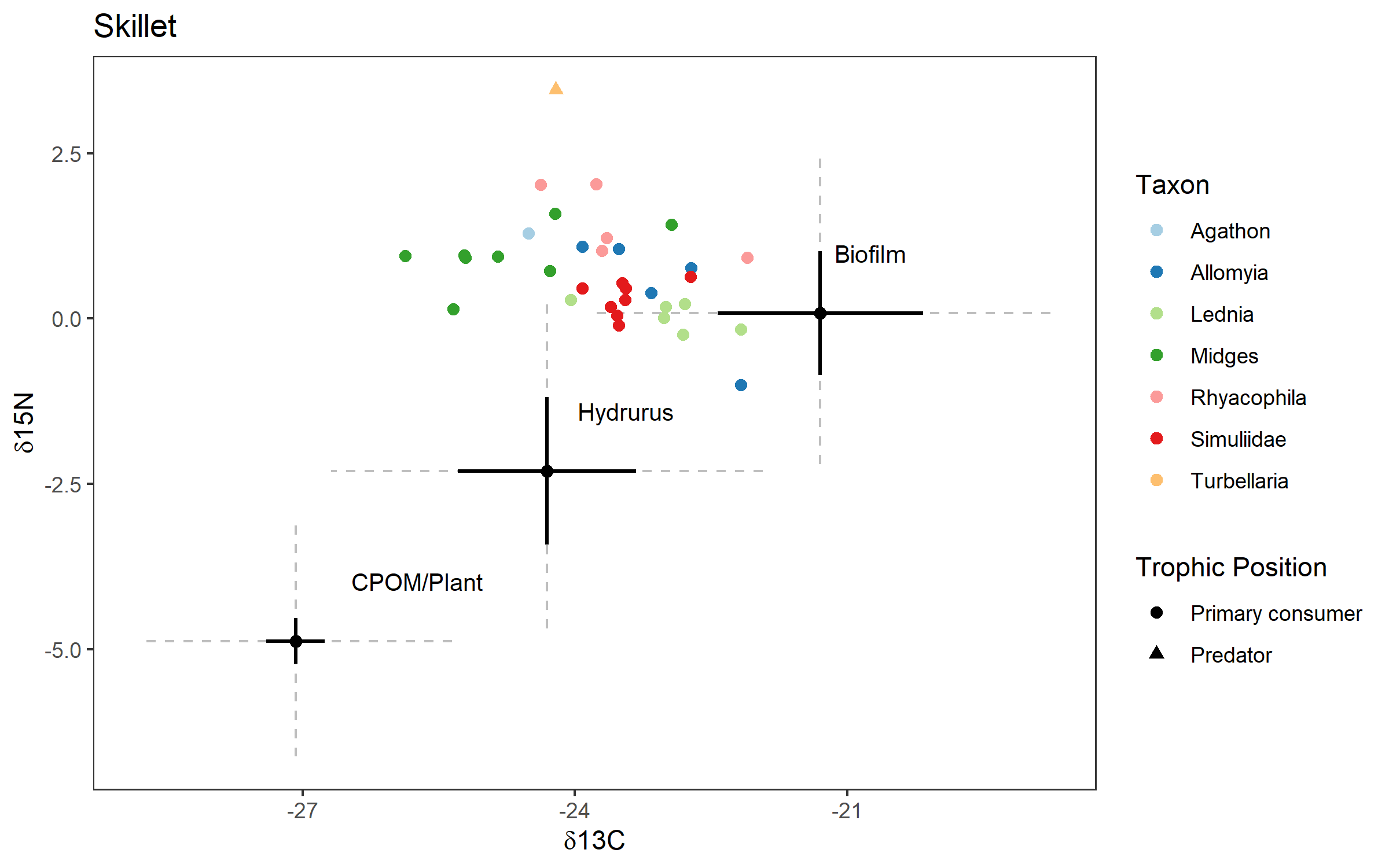

Figure S9. Raw isotope values for Skillet (glacier-fed). Resources are represented as means and standard errors and individual invertebrates are represented as points. Resources were shifted by the TEF (0.4 ‰ ± 1.4 for δ^13^C and 1.4 ± 1.4 for δ^15^N), and grey dashed lines indicate the additional error added by the TEFs.

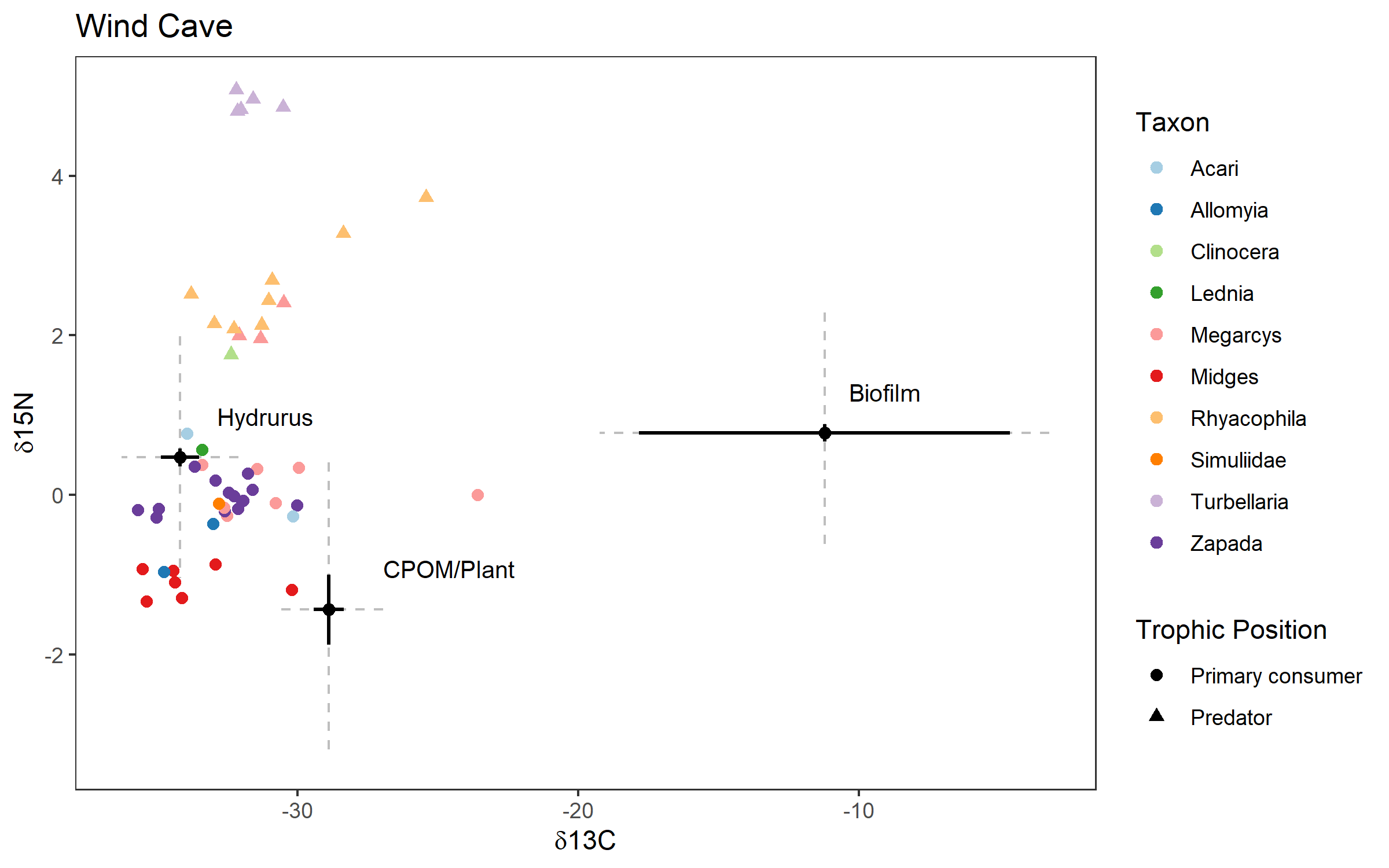

Figure S10. Raw isotope values for Wind Cave (subterranean ice-fed). Resources are represented as means and standard errors and individual invertebrates are represented as points. Invertebrates were shifted by the TEF (0.4 ‰ for δ^13^C and 1.4 for δ^15^N) but source errors represent raw sample error and were not increased to incorporate the TEF uncertainty.

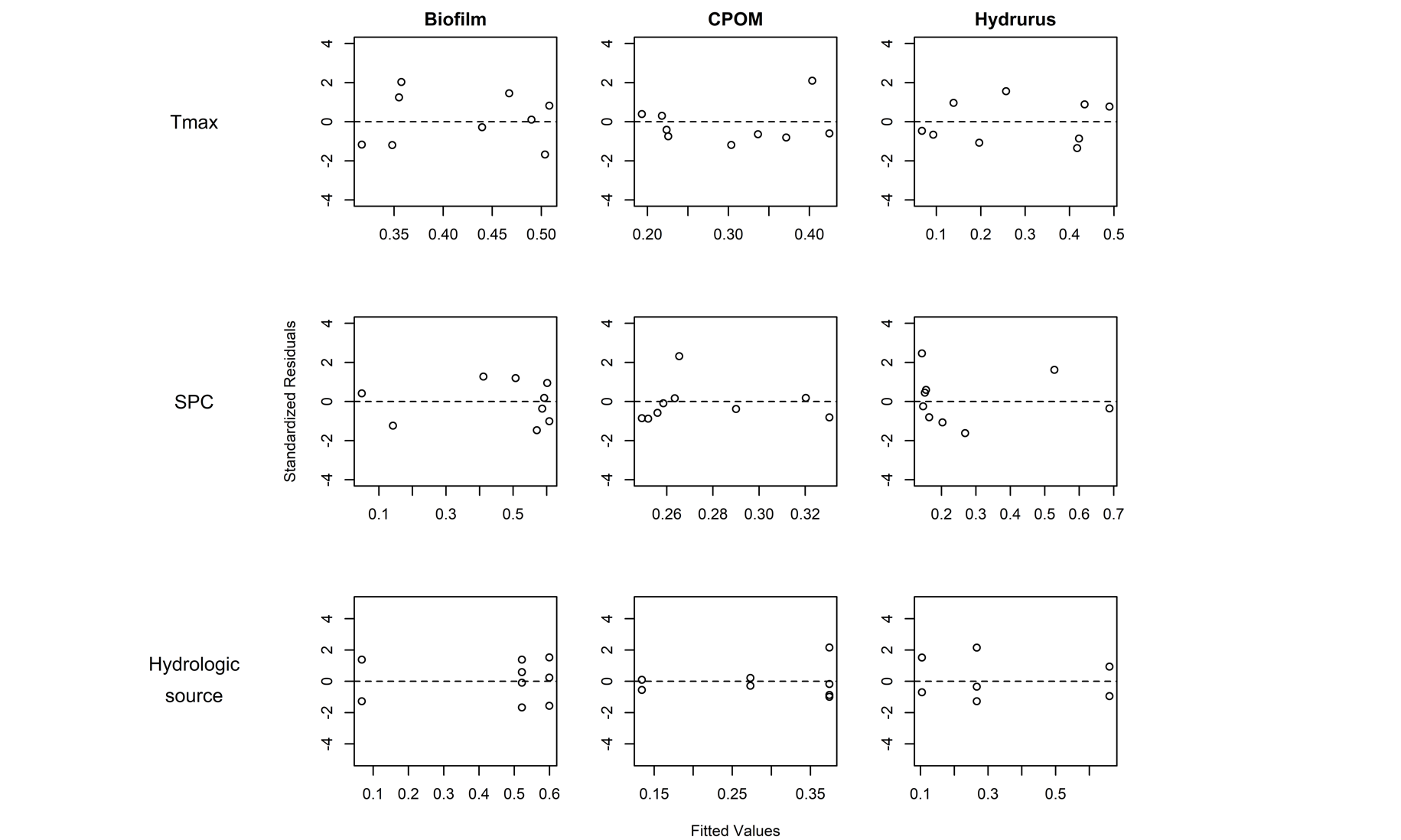

Figure S11. Residuals versus fitted values for maximum temperature (Tmax), specific conductivity (SPC), and hydrologic source models.

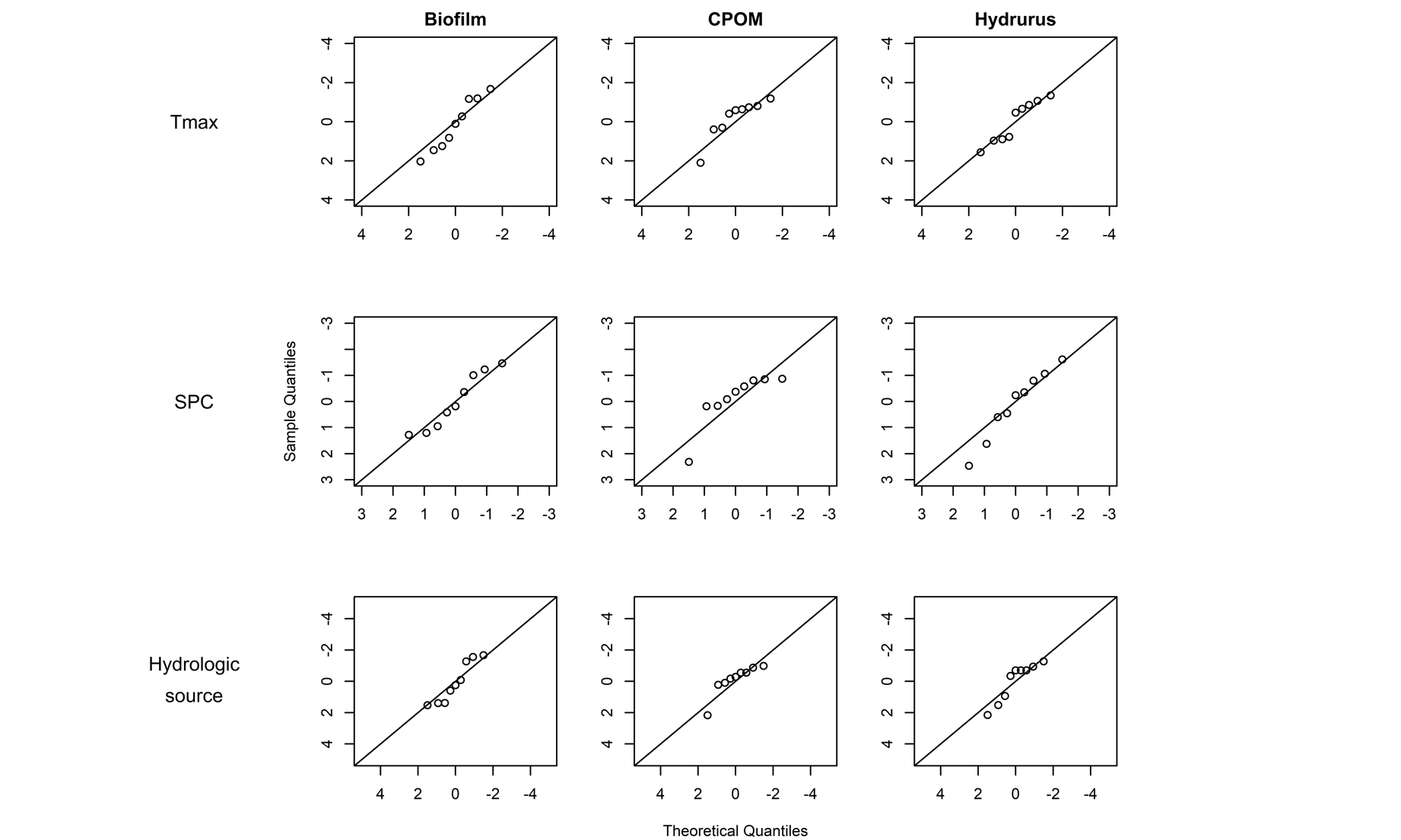

Figure S12. Normal Q-Q plots for specific conductivity (SPC), maximum temperature (Tmax), and hydrologic source models.
